# Systematic analysis of key parameters for genomics-based real-time detection and tracking of multidrug-resistant bacteria

**DOI:** 10.1101/2020.09.24.310821

**Authors:** Claire L Gorrie, Anders Goncalves Da Silva, Danielle J Ingle, Charlie Higgs, Torsten Seemann, Timothy P Stinear, Deborah A Williamson, Jason C Kwong, M Lindsay Grayson, Norelle L Sherry, Benjamin P Howden

## Abstract

**Background:** Pairwise single nucleotide polymorphisms (SNPs) are a cornerstone for genomic approaches to multidrug-resistant organisms (MDROs) transmission inference in hospitals. However, the impact of key analysis parameters on these inferences has not been systematically analysed.

**Methods:** We conducted a multi-hospital 15-month prospective study, sequencing 1537 MDRO genomes for comparison; methicillin-resistant *Staphylococcus aureus*, vancomycin-resistant *Enterococcus faecium*, and extended-spectrum beta-lactamase-producing *Escherichia coli* and *Klebsiella pneumoniae*. We systematically assessed the impact of sample and reference genome diversity, masking of prophage and regions of recombination, cumulative genome analysis compared to a three-month sliding-window, and the comparative effects each of these had when applying a SNP threshold for inferring likely transmission (≤15 SNPs for *S. aureus*, ≤25 for other species).

**Findings:** Across the species, using a reference genome of the same sequence type provided a greater degree of pairwise SNP resolution, compared to species and outgroup-reference alignments that typically resulted in inflated SNP distances and the possibility of missed transmission events. Omitting prophage regions had minimal impacts, however, omitting recombination regions a highly variable effect, often inflating the number of closely related pairs. Estimating pairwise SNP distances was more consistent using a sliding-window than a cumulative approach.

**Interpretation:** The use of a closely-related reference genome, without masking of prophage or recombination regions, and a sliding-window for isolate inclusion is best for accurate and consistent MDRO transmission inference. The increased stability and resolution provided by these approaches means SNP thresholds for putative transmission inference can be more reliably applied among diverse MDROs.

**Funding:** This work was supported by the Melbourne Genomics Health Alliance (funded by the State Government of Victoria, Department of Health and Human Services, and the ten member organizations); an National Health and Medical Research Council (Australia) Partnership grant (GNT1149991) and individual grants from National Health and Medical Research Council (Australia) to NLS (GNT1093468), JCK (GNT1008549) and BPH (GNT1105905).

## BACKGROUND

Antimicrobial-resistant (AMR) pathogens are amongst the foremost threats to global public health ^1–5^. On an individual level, they lead to increased morbidity and mortality, both in terms of the initial infection and resulting sequelae or complications, as well as significant increases in treatment costs and length of hospital stay ^6–8^. Consequently, this places an increasing strain on healthcare systems as the global burden of AMR pathogens rises ^6,8–10^. Among the pathogens of particular concern are multidrug-resistant organism (MDRO) species such as the ESKAPE pathogens (*Enterococcus faecium, Staphylococcus aureus, Klebsiella pneumoniae, Acinetobacter baumanii, Pseudomonas aeruginosa*, and *Enterobacter aerogenes*), and *Escherichia coli* ^1,11–13^.

The World Health Organisation recently highlighted the need to invest in resources to enhance the surveillance of AMR ^3^, which can be facilitated through genomics ^5,14^. Although whole genome sequencing (WGS) is increasingly leveraged in public health outbreak investigations, including for AMR, these have predominantly focused on retrospective ‘closed’ datasets. In these reports, study-specific analysis approaches have defined single nucleotide polymorphism (SNP) thresholds for ruling isolates as ‘likely’ or ‘unlikely’ part of transmission events, based on a combination of genomic and epidemiologic evidence ^15–18^, and some have determined thresholds of genomic diversity between sequences that is correlated with epidemiological transmission evidence (e.g. SNP distance) ^15,16,18^. Whilst these SNP thresholds perform well in a closed dataset, their application to prospective genomic surveillance datasets, with different analysis approaches, needs to be evaluated and developed further, especially when dealing with more complex - genetically or temporally diverse - datasets.

Many of the MDROs posing the greatest health threats exhibit significant population genomic diversity, prolonged asymptomatic colonisation, horizontal gene transfer, and DNA acquired via homologous recombination. These factors can impact relative genetic relatedness, so the methods and transmission SNP thresholds used must remain robust amongst such genomic dynamism. There still remains a significant gap between bespoke comparative genomics research approaches applied in retrospective studies, and the effective translation of such approaches into real-time surveillance in clinical settings in order to help inform infection prevention and control of MDROs.

To address this knowledge gap, we investigated three major facets of genomics data analysis with potential for significant impacts on the accurate surveillance and transmission detection, using a comprehensive genomic and epidemiological dataset for four major hospital MDROs. These were: i) reference genome choice and level of analysis, i.e. species versus sequence type; ii) omission of DNA regions predicted to be prophage or acquired by recombination, and; iii) genome inclusion or exclusion in a growing dataset (cumulative versus a sliding-window approach). We show that the best approach was using a closely related reference genome, without omitting prophage or recombination regions, and a sliding-window for sample inclusion. These methods provided finer-scale resolution and greater consistency and accuracy in pairwise SNP distances for inferring isolate relatedness, making the application of a single SNP threshold to define transmission more appropriate than other approaches. These findings provide the basis for a framework for pathogen-specific standardisation for MDRO surveillance using genomics.

## METHODS

### Isolate selection and whole genome sequencing

During a 15-month prospective study (April – June 2017^19^ and October 2017 – November 2018) all positive clinical or screening samples for four dominant healthcare-associated MDROs were collected for WGS from eight hospitals in Melbourne, Australia. This included all methicillin-resistant *Staphylococcus aureus*, all *vanA* vancomycin-resistant *Enterococcus faecium*, all extended-spectrum beta-lactamase (ESBL) phenotype *Klebsiella pneumoniae*, and all ESBL ciprofloxacin-resistant *Escherichia coli* (in the first eight weeks, ESBL ciprofloxacin-susceptible *E. coli* were also included).

Additional detail on study design, sample collection and identification, and laboratory and sequencing/bioinformatics workflows available in **Supplementary Methods**.

To capture diversity within each species and to focus on the dominant genotypes we selected all sequences representing the four most common multi locus sequence types (STs) of each species (n=153) (**Table 1, Supplementary Table 1**). Short read sequence data available at BioProject PRJNA565795.

**Table 1.** Summary of species, sequence types, and reference genomes for 1537 genomes included in this study.

| Species (n total isolates) | Sequence Type | Number of isolates | Reference chromosome | Reference chromosome size |
| --- | --- | --- | --- | --- |
| *Staphylococcus aureus* (n = 510) | ST5 | 61 | BPH2819 | 2733461 bp |
|  | ST22* | 222 | BPH2900* | 2823339 bp |
|  | ST45 | 158 | NC_021554.1 | 2850503 bp |
|  | ST93 | 69 | NC_017338.1 | 2811435 bp |
| *Enterococcus faecium* (n = 305) | ST80 | 29 | CP027501 | 2912017 bp |
|  | ST203 | 60 | CP027517 | 2863087 bp |
|  | ST1421* | 146 | CP027497* | 2883877 bp |
|  | ST1424 | 70 | AUSMDU00011555 | 2946167 bp |
| *Klebsiella pneumoniae* (n = 62) | ST15 | 12 | CP034045 | 5319653 bp |
|  | ST17 | 12 | CP009461 | 5118878 bp |
|  | ST307* | 23 | CP025146* | 5383248 bp |
|  | ST323 | 15 | CP024499 | 5234963 bp |
| *Escherichia coli* (n = 660) | ST38 | 39 | CP026723 | 5492922 bp |
|  | ST131* | 460 | NC_013654.1* | 4717338 bp |
|  | ST648 | 51 | CP023258 | 5074278 bp |
|  | ST1193 | 110 | CP030111 | 4939457 bp |

\* indicates the reference chromosome in both the species-level (multiple-ST) alignment and the outgroup-reference alignment.

### Mapping and single nucleotide polymorphism (SNP) calling

All mapping and SNP calling analyses were conducted using snippy (v4.6.0, https://github.com/tseemann/snippy, *minfrac* 10 and *mincov* 0·9). Additional detail on all mapping analyses available in **Supplementary Methods**.

### Pairwise SNP distances and transmission inference thresholds

Pairwise SNPs were calculated in R using harrietr (v0.2.3, https://github.com/andersgs/harrietr) and the core SNP alignments. Transmission inference thresholds (≤15 SNPs for MRSA, ≤25 SNPs for other species) were applied. More detail available in **Supplementary Methods**.

### Figures, data visualisation

All figures were created in R (v3.6.0 as above), using one or more of the following packages: ggplot2 (v3.3.1), patchwork (v1.0.0, https://github.com/thomasp85/patchwork), IRanges (v2.18.3, https://github.com/Bioconductor/IRanges), tidyverse (v1.3.0, https://www.tidyverse.org), and RColorBrewer (v1.1-2, https://CRAN.R-project.org/package=RColorBrewer).

### Statistical analyses

All statistical analyses were conducted in R, with more detail available in **Supplementary Methods**.

## RESULTS

### Choice of reference genome, sample size, and population diversity all impact number of SNPs detected

Three different alignment approaches were undertaken for all sequence types (STs) in each species, in order to investigate the impact of reference genome relatedness and isolate diversity. The first was the ‘species alignment’, with all isolates from each species’ four most common STs aligned to the ‘species reference’ (reference chromosome of the same ST as the largest ST for that species and show by * in Table 1). The second alignment for each ST used only isolates of the given ST, but still used the ‘species reference’, herein referred to as the ‘outgroup-reference alignment’. The third alignment was the ‘ST alignment’, using only isolates of any given ST and a reference genome of the same ST. For the most common ST in each species, the species reference was of the same ST, hence the outgroup-reference and ST alignments were identical. We chose to focus on ST as a means to triage and group within species, as it is widely recognised in both the genomic and clinical microbiology fields. Details on the resulting alignments are provided in **Supplementary Table 2**. Phylogenetic trees, including relative position of the reference and population structure, are shown in **Supplementary Figures 1-4**.

Independent of species, 11/16 ST-grouped analyses showed significant differences in distribution of pairwise SNPs distances when comparing the different alignment approaches (**Table 2**, **Figure 1**), indicating that reference genome selection is critical for robust pairwise SNP comparisons. *Enterococcus faecium* ST80 and *K. pneumoniae* ST17 were exceptions, both showing no significant difference between the outgroup-reference alignment and the ST alignment, likely explained by high intra-ST diversity compared to others. *E. faecium* ST80 forms numerous clusters throughout the species phylogeny and *K. pneumoniae* ST17 shows much deeper branching and genetic distance than other STs in the tree (**Supplementary Figures 2, 3**).

**Table 2.**
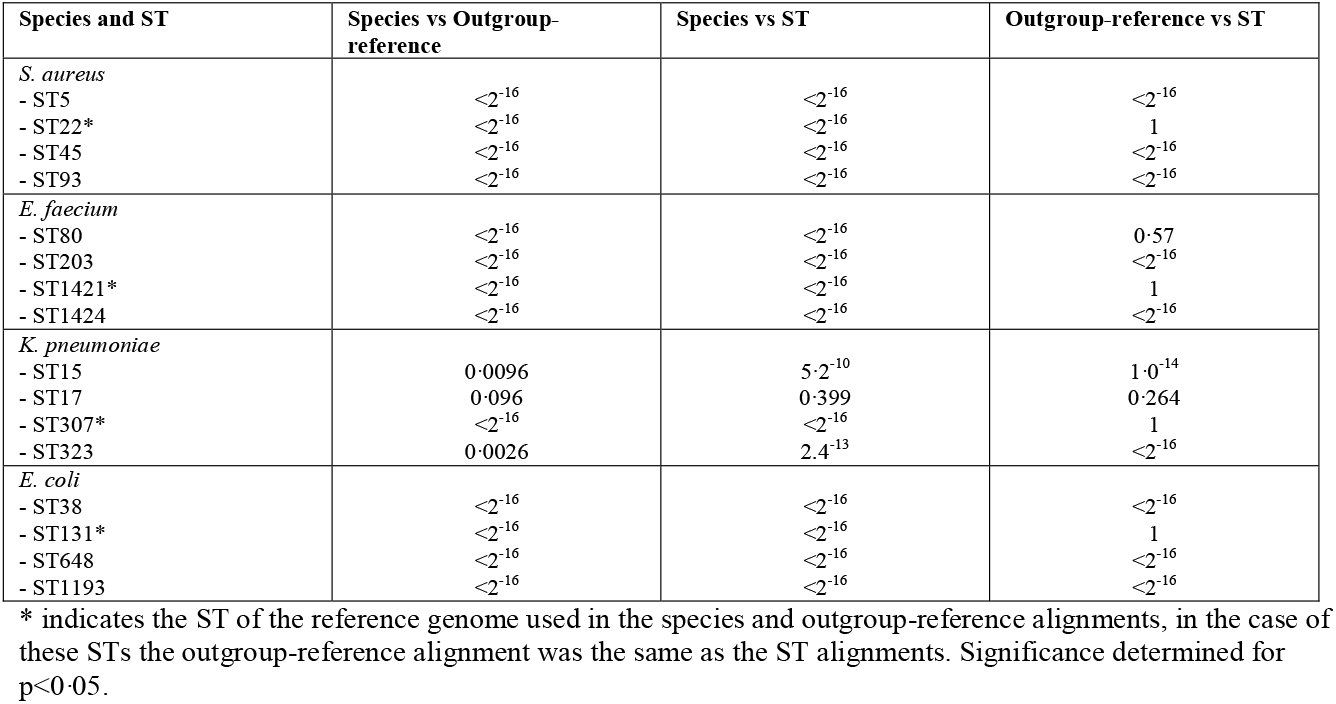
P-values arising from pairwise Wilcoxon tests for significance between species, outgroup-reference and ST alignments for each ST.

**Figure 1.**
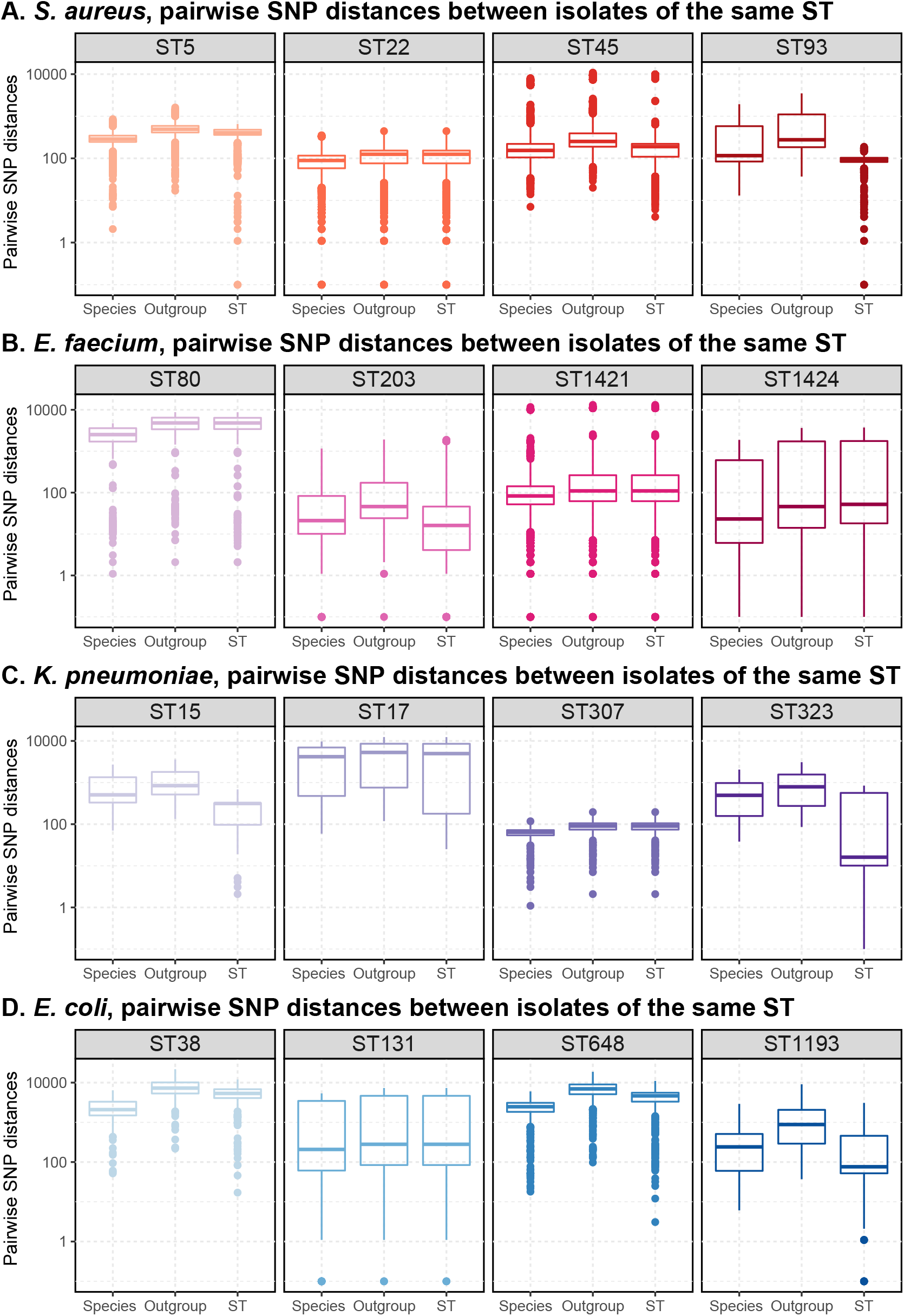
Distribution of single nucleotide polymorphism (SNP) distances between isolate pairs of the same sequence type, from three different reference-alignment combinations. Pairwise SNP distances are shown on log10 scale on the y-axis; maximum y-axis values differ by species. The three reference-alignment comparisons are shown on the x-axis. ‘Species’ shows pairwise SNP distances drawn from an alignment of isolates from four different STs against the species reference genome, as per **Table 1**. This same reference genome is used as an outgroup-reference, shown here under ‘Outgroup’, but all isolates are of a single ST. ‘ST’ uses both isolates and reference genome of the same ST. All boxplots are coloured according to ST.

These analyses demonstrate that, for the majority of species/STs tested, where less genomic diversity is present, the various approaches generate consistently different pairwise SNPs distances. In particular, resolution is lost when using a distant reference resulting in a smaller core alignment and typically higher numbers of pairwise SNP distances; truly closely related isolate pairs may be misclassified as unlikely transmission. In contrast, for highly diverse STs, it can make little difference whether a close or distant reference genome is used.

### Effects of masking prophage and recombination regions

Having established that the ST alignments are generally better for fine-scale analyses, these were used to test the effect of masking regions of horizontal gene transfer. Previous studies have suggested that these regions result in elevated SNP counts meaning that inferred phylogenies do not represent the vertical evolution of the population, which may interfere with identifying transmission through evolution ^20–23^. Regions predicted to be prophage and/or homologous recombination were masked and the resulting pairwise SNP distances compared to those without masking (**Figure 2**).

**Figure 2.**
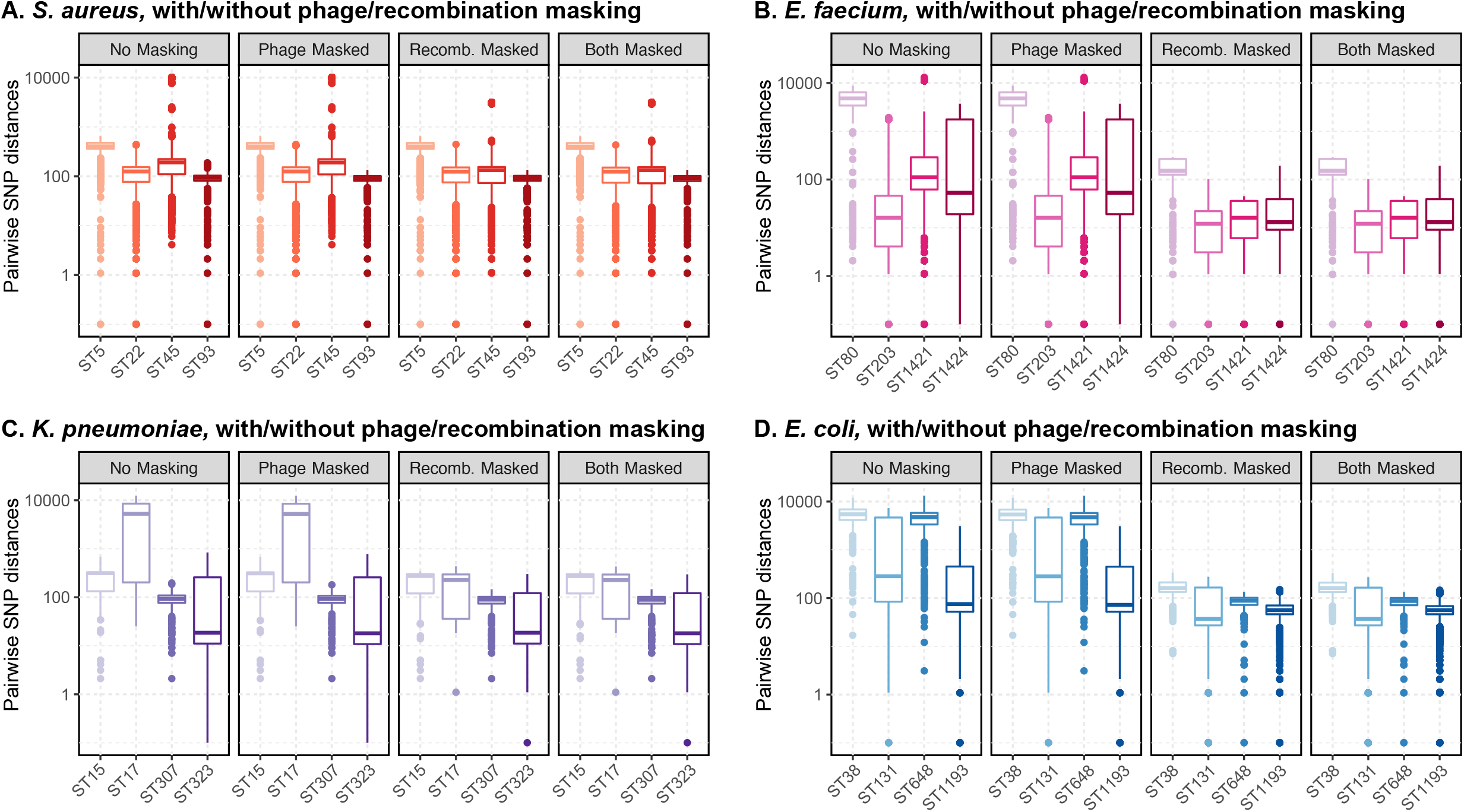
Distribution of single nucleotide polymorphism (SNP) distances between isolate pairs of the same sequence type, before and after masking regions of phage, recombination, or both (phage and recombination). Pairwise SNP distances are shown on log10 scale on the y-axis; maximum y-axis values differ by species. Sequence type (ST) are shown on the x-axis, and boxplots are also coloured by ST.

Across all species, masking prophage regions had little-to-no effect on the core alignment, the core SNP alignment, or pairwise SNP distances (**Figure 2, Supplementary Table 3**). Prophage regions often coincided with regions that were already excluded from analysis as they did not form part of the core genome (as shown in **Supplementary Figures 5-8**). In contrast, recombination masking showed considerable effects, though the effect size differed amongst the various species and STs (**Figure 2, Supplementary Table 3**). The largest differences were among multiple *E. faecium* and *E. coli* STs and *K. pneumoniae* ST17, where recombination masking saw many isolates’ pairwise SNP distances fall by hundreds or even thousands of SNPs. The extent of effect caused by recombination masking clearly correlated with the number and size of regions of recombination (**Figure 3**). For example, some *S. aureus* and *K. pneumoniae* STs (ST5/ST22/ST93 and ST15/ST307/ST323 respectively) each had only a few small recombination regions detected (**Supplementary Figures 5, 7**), and pairwise SNP distances showed minimal changes when this recombination was omitted (**Figure 2A, 2C**), whereas many of the other STs and species had large areas of genome removed due to recombination masking. In the most extreme cases, recombination masking resulted in significant portions of the genome being masked and the average pairwise SNP distances dropping from many thousands of SNPs to hundreds (*E. faecium* ST80 [**Supplementary Figure 6A**] and *K. pneumoniae* ST17 [**Supplementary Figure 7B**]).

**Figure 3.**
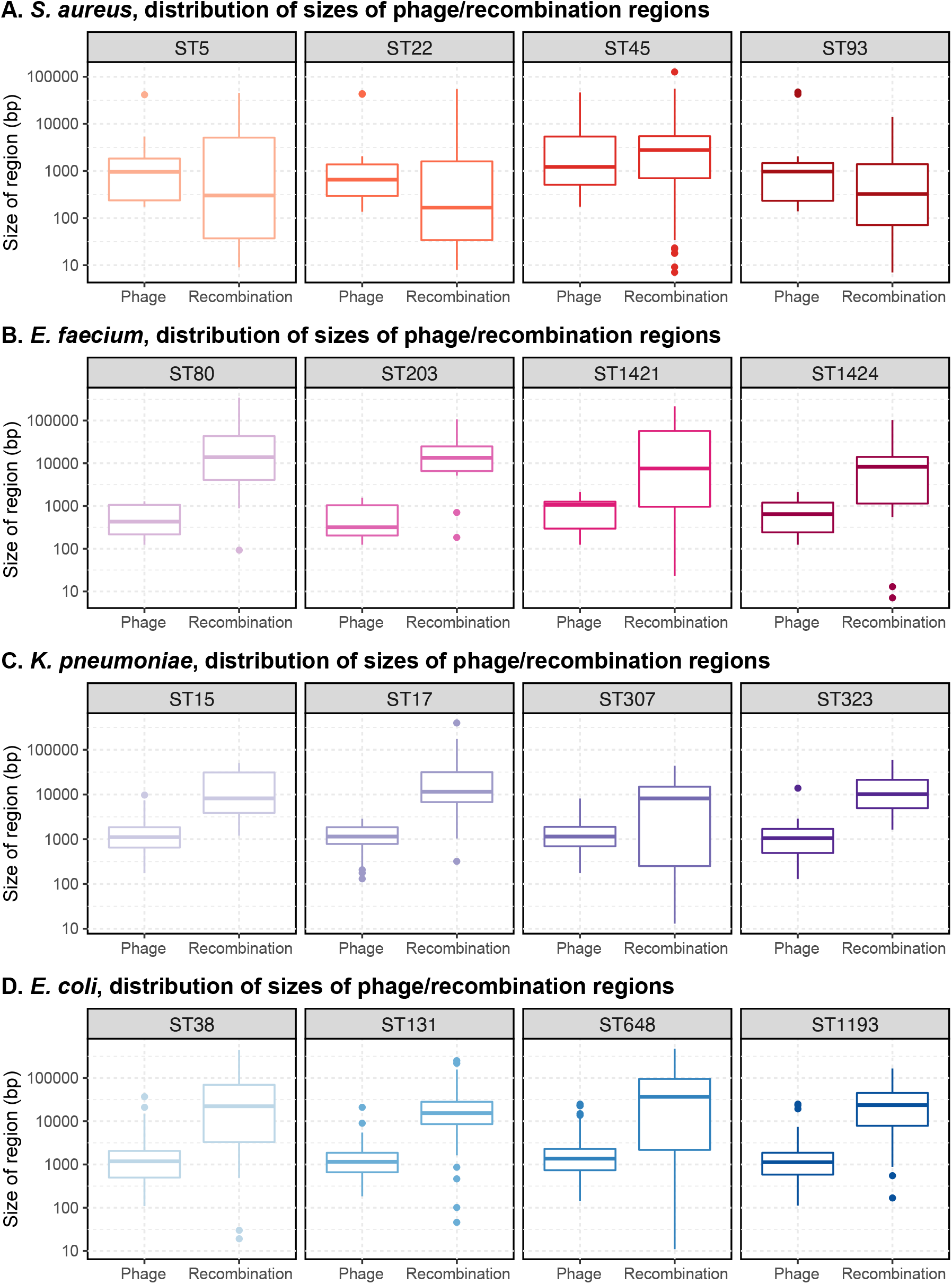
Distribution of predicted phage and recombination region sizes. The size of the region (in base pairs [bp]) is shown on a log10 scale on the y-axis; maximum y-axis values differ by species. The type of region, either phage or recombination, is shown on the x-axis. Boxplots are colour by sequence type (ST).

The combined masking of both prophage and recombination showed very similar results to those seen when masking for only recombination (shown in **Figure 5**); in most species and STs, predicted regions of recombination included those regions that had been predicted to be prophage (as seen in **Supplementary Figures 5-8**).

In cases where isolates are already closely related, masking prophage and/or recombination makes minimal, if any, difference in pairwise SNP distances meaning that transmission inference is unaffected. However, isolate pairs that have many pairwise SNP between them, but which have many of these SNPs masked as regions of recombination, can then erroneously appear to be closely related and could incorrectly be inferred as likely transmission. In these cases, it would be inappropriate and misleading to mask recombination when inferring transmission.

### Effect of cumulative and sliding-window approaches on prospective/real time transmission surveillance and inference

Using the ST alignments, and without masking for prophage or recombination, two different approaches for isolate inclusion and comparison over time were implemented; a cumulative approach where all additional isolates were included over time, and a three-month sliding-window approach. In some cases, isolates potentially arising from the same outbreak are collected over long time periods, and may be important for context and transmission inference. This has been well described for a number of MDRO outbreaks such as drug-resistant *K. pneumoniae* where epidemiologically linked samples have been found over years, in part driven by long-term asymptomatic colonisation ^24^. As such, it is important to establish the potential impact of a continually growing and diversifying dataset, as compared to closed short-term datasets.

In the cumulative approach, all new isolates from each sampling month were compared to all previously included isolates. As the total number of isolates increased over time, so did the diversity, resulting in a continually diminishing core genome alignment (variant and invariant sites) (**Figure 4, Supplementary Table 4**). On average, 17.6% (range: 4%-57%) of the reference genome length was lost from the core alignment from the first to last month of sampling (**Supplementary Table 2**). *E. coli* ST131 had the greatest loss falling from 91% of the reference genome in the first sampling month to only 34% in the final month. The core SNP alignment in most STs increased over time; although the core genome was shrinking, more of the core sites became variant (i.e. SNPs) (**Figure 4, Supplementary Table 4**). *E. coli* ST131 was an exception, with a steady decrease detected in both the core genome and core SNP alignments (**Figure 4**).

**Figure 4.**
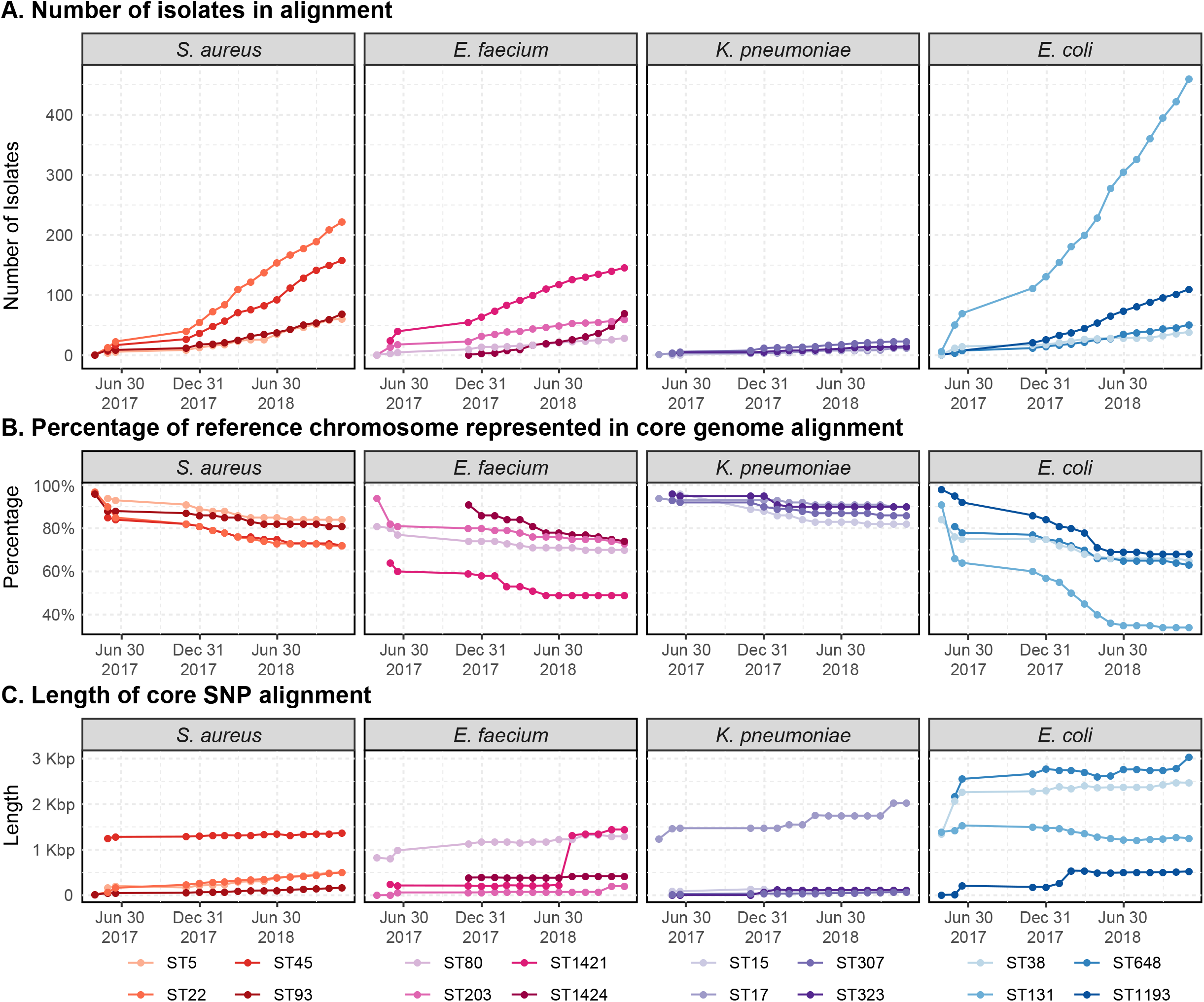
Effects of cumulative inclusion of all isolates over time, calculated at the conclusion of each calendar month. **Panel A:** the total number of isolates collected and included in the alignment and analysis. **Panel B:** the proportion of the reference chromosome that is represented in the core genome alignment (both variant [including SNPs] and invariant sites) as a percentage of the full reference chromosome length. **Panel C:** the length of the core SNP alignment, shown on the y-axis in kilobase pairs (Kbp). All plots are coloured by sequence type (ST).

**Figure 5.**
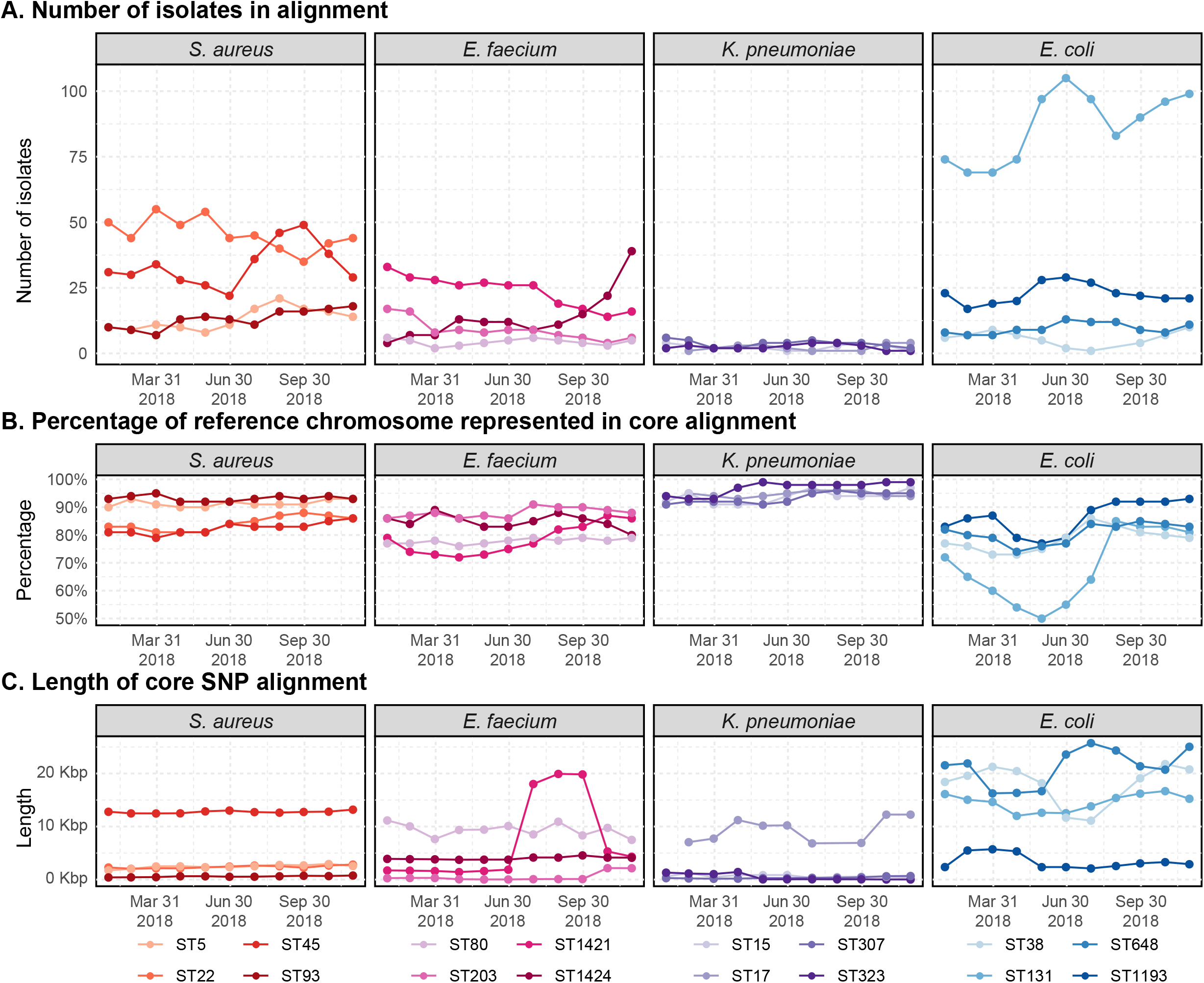
Effects of sliding-window inclusion of isolates over time, calculated at the conclusion of each three-month window. **Panel A:** the total number of isolates collected and included in the alignment and analysis. **Panel B:** the proportion of the reference chromosome that is represented in the core genome alignment (both variant [including SNPs] and invariant sites) as a percentage of the full reference chromosome length. **Panel C:** the length of the core SNP alignment, shown on the y-axis in kilobase pairs (Kbp). All plots are coloured by sequence type (ST).

The sliding-window approach utilised a three-month window, ‘sliding’ forwards by a single month each time. In this approach, although there were fluctuations in the proportion of the reference in the core genome alignment over time, it did not continually decease as with the cumulative approach (**Supplementary Table 5**). The mean core alignment size was consistently higher; more potentially informative sites are present at each time point, providing finer resolution. For example, while the proportion of the reference genome in the core alignment for *E. coli* ST131 was reduced to an average 48% and minimum of 34% in the cumulative approach, the sliding-window approach had an average of 68% and minimum of 50%. In providing much larger and more consistently sized core alignments, the proportion of reference genome represented in the core alignment, it is also easier to compare pairwise SNP distances over time.

### Effect of different approaches on ruling likely or unlikely transmission when applying a SNP distance threshold

Although a SNP threshold is commonly applied to infer likely transmission, the choice of genomic analysis methods has a large influence in calculating the pairwise SNP distances and therefore which isolate pairs fall below the set threshold. Here, we applied SNP thresholds (≤15 SNPs for *S. aureus* and ≤25 SNPs for the other species) to rule isolates as “likely” or “unlikely” putative transmission events for every approach used in this study. We calculated the overall proportion of isolate pairs that fell below the species’ SNP thresholds for likely transmission, and importantly also identified how many pairs were above the SNP threshold for likely transmission in one, or more, of the alignment approaches, but ‘shifted’ below the threshold in another.

When assessing the effect of isolate and reference genome diversity we found that the out-group reference approach provided the lowest number of likely transmission pairs compared to both the species and ST alignments (**Supplementary Table 2, Figure 6**). None of the pairs that experienced a shift below the SNP threshold did so as a result of the outgroup-reference analysis, with the exception of the *E. faecium* ST1424 (**Table 3**). The same was calculated for comparing absence of masking of prophage and/or recombination regions, to the unmasked alignment (details in **Figure 6, Supplementary Table 3**). Again, we calculated the number of pairs shifting below the threshold following masking of any kind (**Table 4**). In almost all cases, masking prophage had little effect on reclassifying isolates pairs to below the SNP thresholds. Conversely, in most species and STs where large amounts of recombination were detected and masked, the number of pairs shifting below the SNP threshold increased by hundreds or, in the case of *E. faecium* ST1421 and *E. coli*. ST131, by thousands. Finally, we considered the effect of the cumulative and sliding-window approaches to sample inclusion (**Figure 6**, **Supplementary Tables, 5**). We identified any ‘shift’ below the SNP threshold observed between the first and last observation of each pair compared ≥2 times (**Table 5**).

**Figure 6.**
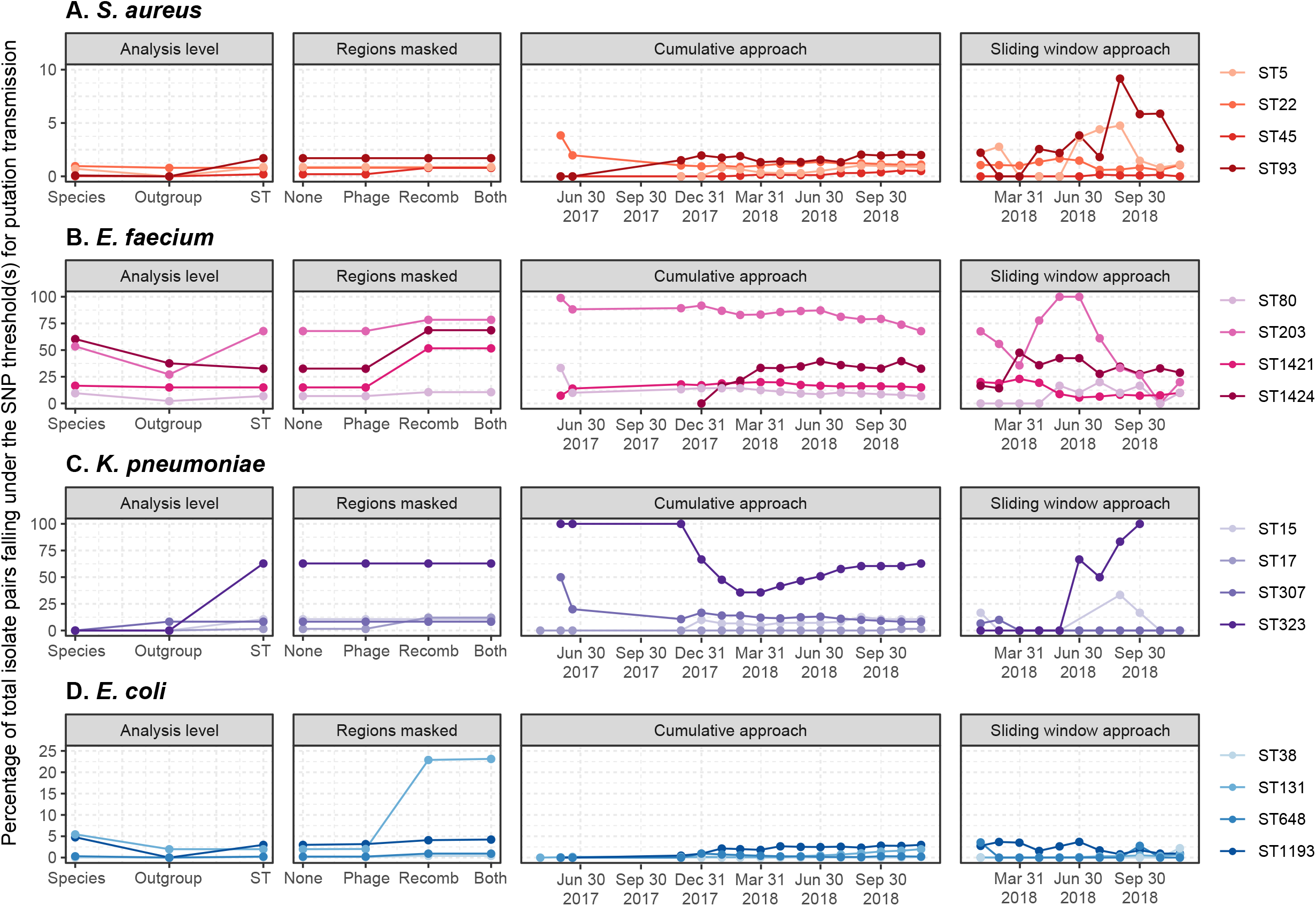
Effects of analysis level, masking of phage and/or recombination regions, and different approaches to sample inclusion of time, on proportion of isolate pairs falling under the SNP threshold for putative transmission. The y-axis shows the percentage of isolates pairs under the SNP threshold for putative transmission for each species; for *S. aureus* (**Panel A**) the threshold is ≤15 SNPs, for all other species (**Panel B-D**) the threshold is ≤25 SNPs. The y-axis maximum value differs by species but is consistent across all plots for the species. All plots are coloured by sequence type (ST).

**Table 3.** Isolate pairs variably below SNP threshold with the three analysis approaches. Total number of pairs that are variably below the threshold for some, but not all, of the analysis approaches, shown for each species and ST, as well as the number of those pairs that experience a shift, that are below the SNP threshold in each of the analysis.

| Species and ST<br>(total isolate pairs) | Total variable pairs | Number of variable pairs below SNP threshold for each analysis |  |  |
| --- | --- | --- | --- | --- |
|  |  | Species | Outgroup | ST |
| *S. aureus* |  |  |  |  |
| - ST5 | 16 | 13 | 0 | 16 |
| - ST22* | 40 | 40 | 0 | 0 |
| - ST45 | 24 | 3 | 0 | 24 |
| - ST93 | 40 | 2 | 0 | 40 |
| *E. faecium* |  |  |  |  |
| - ST80 | 30 | 30 | 0 | 19 |
| - ST203 | 785 | 466 | 0 | 720 |
| - ST1421* | 169 | 169 | 0 | 0 |
| - ST1424 | 670 | 670 | 120 | 0 |
| *K. pneumoniae* |  |  |  |  |
| - ST15 | 7 | 0 | 0 | 7 |
| - ST17 | 1 | 0 | 0 | 1 |
| - ST307* | 3 | 3 | 0 | 0 |
| - ST323 | 66 | 0 | 0 | 66 |
| *E. coli* |  |  |  |  |
| - ST38 | 1 | 0 | 0 | 1 |
| - ST131* | 3678 | 3678 | 0 | 0 |
| - ST648 | 4 | 4 | 0 | 3 |
| - ST1193 | 425 | 286 | 0 | 179 |

\* indicates the sequence of the reference genome used for the species and outgroup analyses.

**Table 4.** Isolate pairs variably below SNP threshold with one or more of the masking approaches. Total number of pairs that are above the SNP threshold without masking but that shift below the threshold with one or more masking approaches, shown for each species and ST. The final four columns show the number of these variable pairs that fall below the SNP thresholds for each masking approach.

| Species and ST | Total variable pairs | Number of variable pairs that are below SNP threshold, with each masking approach |  |  |  |
| --- | --- | --- | --- | --- | --- |
|  |  | None | Phage | Recomb. | Both |
| *S. aureus* |  |  |  |  |  |
| - ST5 | 0 | - | - | - | - |
| - ST22 | 1 | 0 | 1 | 0 | 1 |
| - ST45 | 72 | 0 | 0 | 72 | 72 |
| - ST93 | 0 | - | - | - | - |
| *E. faecium* |  |  |  |  |  |
| - ST80 | 15 | 0 | 0 | 15 | 5 |
| - ST203 | 187 | 0 | 0 | 187 | 187 |
| - ST1421 | 3879 | 0 | 0 | 3879 | 3879 |
| - ST1424 | 870 | 0 | 0 | 870 | 870 |
| *K. pneumoniae* |  |  |  |  |  |
| - ST15 | 0 | - | - | - | - |
| - ST17 | 7 | 0 | 0 | 7 | 7 |
| - ST307 | 0 | - | - | - | - |
| - ST323 | 0 | - | - | - | - |
| *E. coli* |  |  |  |  |  |
| - ST38 | 2 | 0 | 0 | 2 | 2 |
| - ST131 | 22339 | 0 | 49 | 22083 | 22339 |
| - ST648 | 9 | 0 | 0 | 9 | 9 |
| - ST1193 | 75 | 0 | 11 | 66 | 75 |

Phage; masking of prophage regions. Recomb; masking of recombination regions.

**Table 5.**
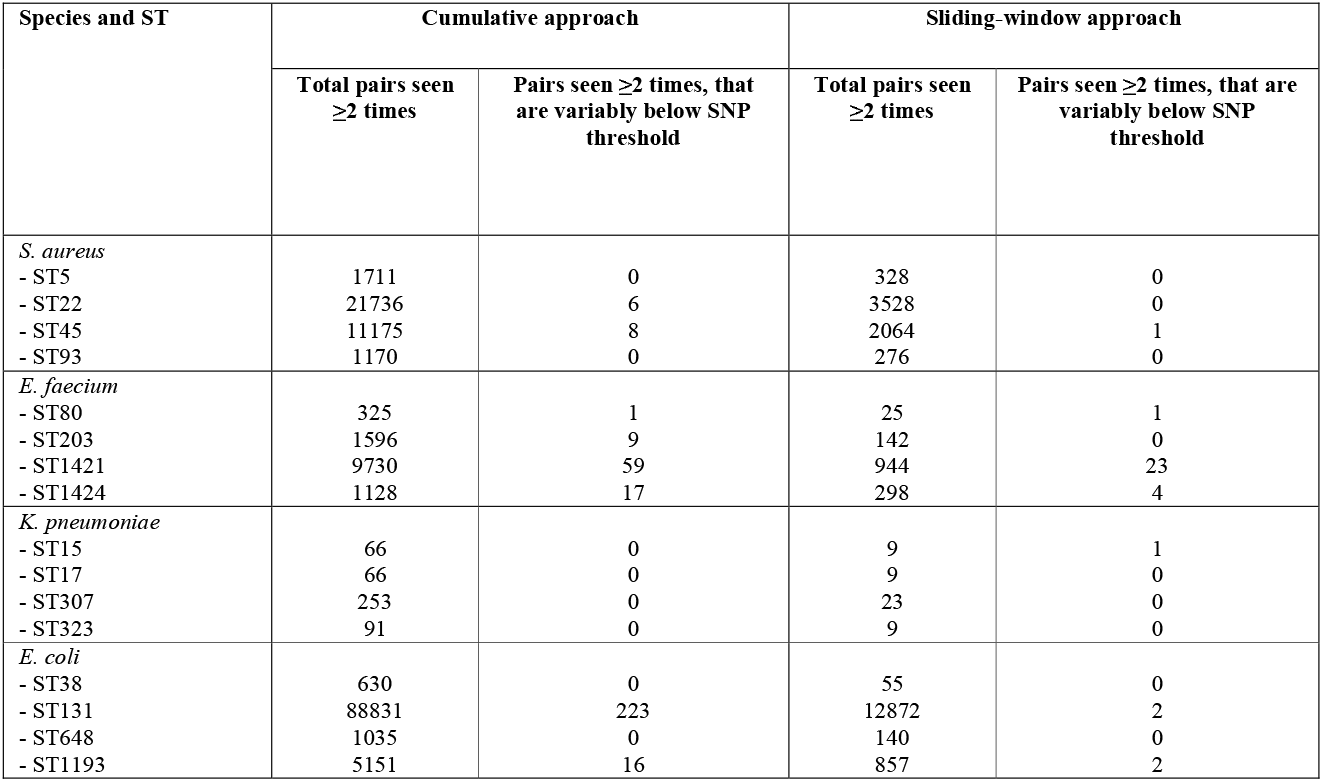
Isolate pairs variably below SNP threshold with either of the isolate inclusion approaches. Total number of variable pairs is shown for each species and ST. In the cumulative approach when a shift below the threshold occurred is was also a shift downwards over time. In the sliding-window approach, the shift could either move from above to below the threshold, or the reverse.

## DISCUSSION

Prospective WGS of hospital MDROs will enhance real-time transmission identification, leading to optimised infection prevention and control and limiting further spread, however methods need to be standardised. Previous studies are often retrospective and *ad hoc*, frequently tailored to a specific, narrow dataset, such as closely related isolates from a single pathogen sequence type or a rare AMR phenotype ^18,24–26^. This is not the reality of prospective hospital or jurisdictional wide surveillance where multiple pathogens and sequence types are detected over time ^27^. As such the results, methods, and thresholds that have been used are not necessarily broadly applicable for prospective surveillance where the dataset continues to expand over time. Here, we utilised a multi-institutional MDRO dataset to systematically investigate a range of approaches on the outcome of potential transmission analyses, providing recommendations for future implementation (**Figure 7**).

**Figure 7.**
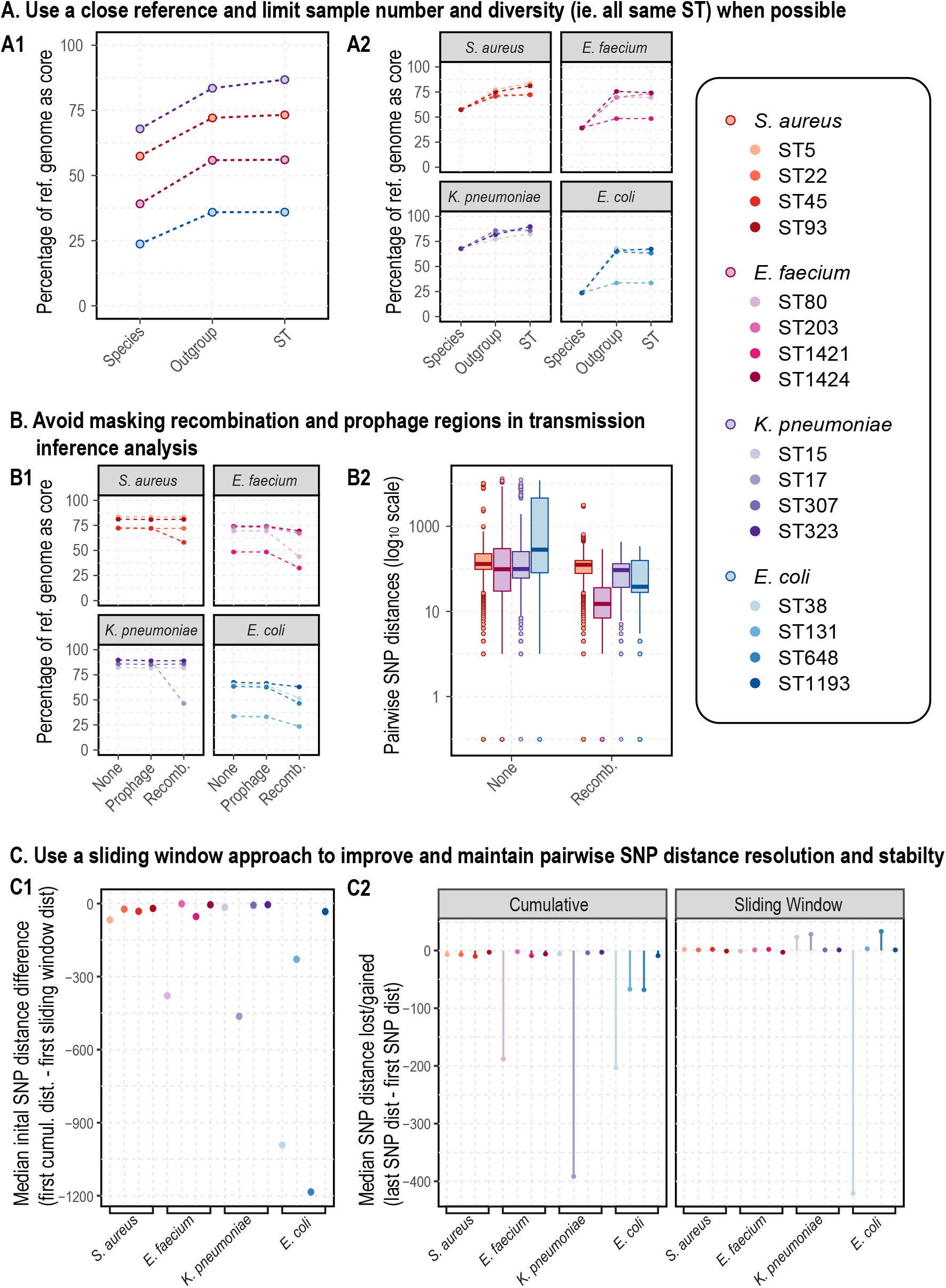
Framework recommendation and justification for pathogen-specific standardisation for MDRO surveillance using genomics. Panel A shows the percentage of the reference genome that is represented in the core genome for each Species (A1) and ST (A2), with each of the three different alignment approaches (shown on the x-axis). Panel B shows the percentage of the reference genome that is represented in the core genome for each ST (B1), with each of the three main approaches to masking regions of horizontal gene transfer (ie. no masking, masking of prophage, and masking of recombination regions; shown on the x-axis). Panel B2 shows the distribution of pairwise SNP distances between all isolate pairs, grouped by Species, without any masking and with masking of recombination regions. Panel C1 shows the median difference between the initial pairwise SNP distances, for all pairs compared in both the cumulative and sliding window approaches that had changing SNP distances observed over time, calculated by subtracting each initial sliding window pairwise SNP distance from each initial cumulative pairwise SNP distance; all values here are less than zero indicating the median initial cumulative pairwise SNP distance is always less than the median initial sliding window pairwise SNP distance. Panel C2 shows the median difference (ie. increase or decrease) between the initial and final pairwise SNP distances, for all pairs compared at least twice, for both the cumulative and sliding window approaches; a negative value shows a median decrease in pairwise SNP distances and therefore a loss of genetic resolution over time as the dataset changes or grows, a positive value shows the opposite. Points and plots are coloured by Species and ST, according to the legend, dotted lines are used (in panels A1, A2, B1) for ease of visualising the relationship between discrete approach variables.

Using a more distant reference genome inflates pairwise SNPs distances, increases ancestral SNPs, and decreases the number of SNPs that have arisen more recently, hence losing the fine-scale resolution required for transmission inference. Although pairwise SNP inflation is lessened when isolates from multiple STs - representing greater genetic diversity and therefore reducing the core genome - are included, this still fails to replicate the pairwise SNP distances seen when using a closer reference. This is generally consistent with previous work ^28^, though for some STs with high genetic diversity and multiple distinct clusters in the phylogenies, it appears to make little difference whether an outgroup reference genome or one of the same ST is used. Given the increased core genome size and fine-scale resolution among more closely related isolate pairs offered when using a closely related reference genome, we recommend doing this wherever possible, though the increased accuracy provided by doing this will be reduced when isolates are highly diverse (**Figure 7, panel A**). Prophage masking had little effect in this dataset primarily due to the fact that the prophage sites corresponded to regions that were already absent from the core genome alignment. In datasets where this is not the case the effect may change but should be assessed. Masking recombination had varying effects, heavily dependent on the individual ST datasets. In cases where isolates were closely related prior to masking, there was little effect, with the opposite seen in more diverse STs. The number and size of recombination regions, as well as the extent of the effect of masking, should be carefully considered; a pair of isolates that have a small number of SNPs after masking but had a hundred regions masked spanning thousands of SNPs, should not be considered as closely related as a pair that had a small number of pairwise SNPs both before and after masking. Masking of prophage and recombination should therefore not be routinely applied for the species discussed here; the former appears to have minimal effects but increases time and effort required, and the latter has the potential to inappropriately reduce the number of SNPs between truly distant isolates (**Figure 7, panel B**). Exceptions may occur when prophage regions are conserved across all isolates or when recombination is limited to a few large regions.

Finally, determining putative transmission often revolves around ruling isolates ‘in’ or ‘out’ of a particular genomic cluster, based on set genomic thresholds and supported by epidemiological analyses. In a truly real-time dataset, new isolates will be continually added over time. The four species in this study can all reside as asymptomatic commensal organisms and can remain undetected for a long time, unless carriage-screening is undertaken, and during this time can undergo diversifying evolution within the host. Given the shrinking core genome and core SNPs, it is also possible that isolate pairs that are distantly related at an initial time point may lose much of that measurable genetic distance by the final timepoint (**Figure 7, panel C2**). This presents at least two serious issues in determining genetic relatedness. If using a threshold to rule transmission in or out, this isolate pair would be initially ruled out and subsequently ruled in as putative transmission. Scaling the SNP numbers proportionate to the amount of core genome or to the entire reference genome may lessen these effects, but if the parts that are lost from the core over time are the more diverse, these scaled or adjusted numbers will still fall short of the true diversity. Though many of these problems are true of the cumulative approach to sample inclusion, they are minimised when using a sliding-window approach. Core genome and SNP alignments, and relative pairwise SNP distances, remain more stable over time, making it easier to standardise or draw comparisons between isolate pairs over time. This approach is also less computationally intensive, given the smaller number of isolates at each time point. It should be noted that it is possible that links between closely related isolates may be missed with the sliding-window approach, if genetically close isolates are temporally more distant. However, using approaches such as single-linkage methods (to identify relatedness between windows) may be used to remedy this, in order to highlight ongoing or persistent transmission chains. For example in time period one, isolates ‘A’ and ‘B’ are closely related and in time period two, isolates ‘B’ and ‘C’ are closely related in the second time period, although isolate ‘A’ is not within time period two we can infer that although separated by time, ‘A’ is related to ‘C’ through ‘B’.

Ultimately, in the context of determining putative transmission it is likely that a SNP threshold will be implemented to rule isolates ‘in’ or ‘out’ on transmission events. However, whilst the threshold may be set, we have demonstrated changes in analysis or the addition of isolates time can see isolate pairs shifting from above the SNP threshold, and therefore ruled ‘out’, to below the threshold and subsequently ruled ‘in’. The most dramatic influence here, in terms number of whether a given pair sat above or below the threshold, were when masking regions of recombination, followed by using more or less distant reference genomes and more diverse (multi-ST) datasets in the alignment. Interestingly, despite the shrinking core alignments observed over time in the cumulative isolate inclusion approach, compared to the sliding-window approach, we saw relatively small numbers of isolates switching from above to below the SNP threshold. However, a larger influence was seen among the more genetically diverse STs (*E. faecium* ST1421 and the *E. coli* ST131).

In summation, when implementing WGS for transmission surveillance of common MDROs we recommend using a closely related genome, without masking of prophage or recombination regions, and a sliding-window approach (**Figure 7**). These all contribute to maximising the SNP distance resolution and stability in an evolving, real-time dataset, and these findings help fill the knowledge gap that has hindered the effective implementation of real-time genomic MDRO surveillance in clinical settings.

## LIST OF ABBREVIATIONS

AMR: Antimicrobial resistant
ESBL: Extended-spectrum beta-lactamase
Mbp / Kbp / bp: Mega-base pair / Kilo-base pair / base pair
MDR: Multidrug-resistant
MDRO: Multidrug-resistant organism
MLST: Multi-locus sequence type
SNP(s): Single nucleotide polymorphism(s)
ST: Sequence type
WGS: Whole genome sequencing

## DECLARATIONS

### Ethics approval and consent to participate

This study was approved by the Melbourne Health Human Research Ethics Committee (HREC) and endorsed by the corresponding HREC at each participating site.

### Consent for publication

Not applicable

### Availability of data and materials

Raw sequence data has been uploaded to the Sequence Read Archive under BioProject PRJNA565795.

### Authors’ contributions

BPH and MLG designed and managed the Controlling Superbugs Study. BPH, CLG and NS designed this project. CLG conducted all genomic, bioinformatic and statistical analyses, and produced the manuscript and all accompanying figures and tables. AGDS was part of the Controlling Superbugs Study Group for the initial project and provided guidance/insights and proofread/edited the manuscript full. DJI provided ongoing input and discussion and edited the manuscript at various stages. CH helped with quality control for both sequence and epidemiological data, as well as conducting long read sequencing and assembly of the E. faecium ST1424 reference genome, and proofread/edited the manuscript. TS wrote a python script/code to calculate which sites in the reference genome were categorised as core sites, and provided bioinformatic advice. TPS provided guidance and feedback both during the study and for the final manuscript. JCK was part of the Controlling Superbugs Study Group for the initial project. NLS was part of the Controlling Superbugs Study Group for the initial project, helped with data collection and quality control, provided guidance and input throughout, and edited the manuscript.

## Acknowledgements

The authors would like to acknowledge the other members of the Controlling Superbugs Study Group include Robyn Lee (previously MDU, currently University of Toronto), Rhonda Stuart (Infectious Diseases, Monash Health; Medicine, Monash University), Tony Korman (Infectious Diseases and Microbiology, Monash Health; Medicine, Monash University), Caroline Marshall (VIDS, Melbourne Health; Peter Doherty Institute), Hiu Tat Chan (Microbiology, Melbourne Health), Maryza Graham (Infectious Diseases and Microbiology, Monash Health; Medicine, Monash University), Paul Johnson (Infectious Diseases, Austin Health; Peter Doherty Institute; Medicine, Austin Health), Marcel Leroi (Microbiology, Austin Health), Caroline Reed (Microbiology, Melbourne Health and Peter MacCallum Cancer Centre), Michael Richards (VIDS, Melbourne Health; Peter Doherty Institute), Monica Slavin and Leon Worth (Infectious Diseases, Peter MacCallum Cancer Centre; National Centre for Infections in Cancer; University of Melbourne), Elizabeth Grabsch (Microbiology, Austin Health), Joanna Price and Carolyn Tullett (Infection Control, Austin Health), Despina Kotsanas (Microbiology, Monash Health), Louise Wright (Infection Control, Monash Health), Suraya Hanim Abdullah Hashim and Jennifer Mitchell (Infectious Diseases, Melbourne Health), Tram Nguyen and Margaret Savanyo (Microbiology, Melbourne Health), Olivia Smibert and Victoria Madigan (Infectious Diseases, Peter MacCallum Cancer Centre) and Peter Pham (MDU Public Health Laboratory, University of Melbourne). Additionally, the authors would like to acknowledge the assistance of the following people: John Greenough, Jennifer Breen and Penny Birchmore (infection control queries and patient movement data) and Carol Wedge (data entry).

