## Supplementary Methods, Supplementary Figures and Figure legends, and Supplementary Tables legends for "Systematic analysis of key parameters for genomics-based real-time detection and tracking of multidrug-resistant bacteria"

#### Study design

We performed a prospective multicentre genomics implementation study of multidrug-resistant organism (MDRO) transmission in eight hospitals from four independent hospital networks, including approximately 2800 acute and subacute inpatient beds, in Melbourne, Australia (population 4.9 million in 2018). MDRO isolates from six defined species (see below) were collected from patient samples (either clinical or screening samples) collected routinely from hospital inpatients (>24h) at any time during their admission; no additional isolates were collected for this study. Duplicate screening isolates were excluded, and duplicate clinical isolates were only excluded if collected less than 14 days after the previous sample. The study was conducted in two phases: a pilot phase (24th April – 18th June 2017) [1], and an implementation phase (30th October 2017 – 30th November 2018), totalling 15 months. Date of collection for each sample was also recorded.

#### MDRO definitions

Four MDROs were included in the pilot phase: *vanA* vancomycin-resistant *Enterococcus faecium* (confirmed by PCR), methicillin-resistant *Staphylococcus aureus* (positive cefoxitin screen or oxacillin MIC >2mg/L), and extended-spectrum beta-lactamase phenotype (EBSL) *Escherichia coli* and *Klebsiella pneumoniae* (defined by ceftriaxone resistance with MIC ≥4mg/L; AmpC phenotypes also included). Due to the large volume of ESBL *E. coli* included during the pilot phase, we narrowed inclusion criteria for the implementation phase to only include ESBL *E. coli* that were also resistant to fluoroquinolones (ciprofloxacin MIC≥2mg/L), as this was more consistent with infection control practices at participating sites (patients with ESBL *E. coli* plus resistance to ≥2 additional antibiotic classes were often isolated using contact precautions).

#### Laboratory workflows

(a) Hospital laboratory – MDROs were isolated, worked up and reported by the hospital laboratory as per their usual protocols. For patients and isolates meeting inclusion criteria, a pure subculture was sent to the central laboratory for sequencing and isolate storage. Results from automated susceptibility testing (Vitek 2 platforms, bioMérieux) were collected from each laboratory.

(b) Sequencing laboratory – A single colony from the subculture received from the hospital laboratory was subcultured onto horse blood agar, incubated overnight, then 1-2 colonies were selected and placed into lysis buffer. DNA extraction, library preparation and quality control (QC) were performed as previous described[1], and isolates sequenced on Illumina NextSeq (San Diego, CA, USA) to achieve 150 bp paired-end read sets.

#### Quality control and species confirmation

All sequences and assemblies underwent quality control checks as part of standard laboratory workflow; sequences not meeting predefined quality metrics were re-sequenced (target sequencing depth ≥40X, minimum average quality score 30). Sequences were assembled *de novo* using *Shovill* (v1.0.4, https://github.com/tseemann/shovill). Species identification was confirmed using *k*-mer identification[2]. To identify *Klebsiella* species that are typically indistinguishable from *K. pneumoniae* by MALDI-ToF, sequences were analysed using the K*leborate* tool (v0.3.0, https://github.com/katholt/Kleborate); subspecies other than *K. pneumoniae sensu stricto* were excluded.

#### Multi-locus sequence typing

Multi-locus sequence typing (MLST) was conducted *in silico* with the *mlst* tool (v2.17.6, https://github.com/tseemann/mlst), using either pubMLST database (for *S. aureus, E. faecium,* and *E. coli*) or BIGSdb Institut Pasteur database (for *K. pneumoniae*)[2].

#### Antimicrobial resistance gene detection

Acquired antimicrobial resistance genes were detected using the *abricate* tool (v0.9.5, https://github.com/tseemann/abricate, minimum coverage and gene identity 100%), employing the NCBI Bacterial Antimicrobial Reference Gene Database (database version 2019-02-08)[3]. Sequences from *E. faecium* isolates were checked to ensure presence of a complete *vanA* operon; if this was absent, further phenotypic workup was performed to look for a *vanA*-containing isolate. ESBL *E. coli* and ESBL *K. pneumoniae* sequences were examined for the presence of ESBL/AmpC genes; if these were absent, isolates were re-tested for ceftriaxone resistance, and only included if resistance was confirmed. Sequences from *S. aureus* isolates were checked for presence of *mecA* (no other *mec* types were detected); if this was absent, cefoxitin and/or oxacillin resistance were confirmed before inclusion.

**Mapping, SNP calling, and sample inclusion**

All mapping and SNP calling analyses were conducted using snippy (v4.6.0, https://github.com/tseemann/snippy, *minfrac* 10 and *mincov* 0.9). To investigate the impact of reference genome selection, we examined the number of SNPs between isolates of the same ST from three different alignment approaches: i) a species alignment, including isolates from four different STs and using a 'species reference' (reference genome of the species' most common ST); ii) an outgroup-reference alignment, using isolates from a single ST but a reference of a different ST (the 'species reference'), and; iii) an ST alignment, using only isolates and a reference genome of the same ST. For the analysis of the cumulative or sliding-window approaches for sample inclusion, the ST alignment approach was used, but subsampled isolates for inclusion in each round of mapping and SNP calling, using snippy-core (from snippy v4.6.0). The cumulative approach included all isolates from the first calendar month of collection (May 2017), then added all isolates from each consecutive sampling month; there was a gap of several months between study phases in which no samples were collected. The sliding-window approach used only isolates from the second collection period of the project, starting November 2017, as the samples from the initial collection period were too distant in time to be included in the three-month time span in this approach.

**Pairwise SNP distances and transmission inference thresholds**

Pairwise SNPs were calculated in R using harrietr (v0.2.3, https://github.com/andersgs/harrietr) and the core SNP alignments. Scaled pairwise SNP distances were calculated by dividing the raw pairwise SNPs between isolates pairs, by the length of the core alignment and multiplying this by either the full length of the reference genome chromosome or a length of 1 Mbp. SNP thresholds were defined based on previous literature and applied to determine impacts of analysis approaches on the inclusion/exclusion of isolates. Isolates with pairwise SNP distances falling below this threshold were determined to be closely related genomically and therefore within the bounds of putative transmission. The SNP thresholds used were ≤15 SNPs for *S. aureus*[3-5] and ≤25 SNPs for *E. faecium*[6,7], *K. pneumoniae* and *E. coli*[8-12].

**Phylogenetic trees**

All phylogenetic trees were inferred using IQtree (v1.6.12)[13] with constant sites, 1000 bootstraps, and a generalised time-reversible model of evolution (GTR+G4).
All trees were midpoint rooted and visualised in R (v3.6.0, https://www.R-project.org/)[14] using ape (v5.4)[15], ggtree (v1.16.6)[16] and ggplot2 (v3.3.1)[17].

**Prophage and recombination region prediction and masking**

All reference genomes were screened using phastaf (v0.1.0, https://github.com/tseemann/phastaf) and the PHASTER database[18,19]. The full dataset for each ST was mapped to the reference of the same ST and Gubbins[20] was used to predict regions of recombination, which were then masked using the masking option in snippy.

**Statistical analyses**

All statistical analyses were conducted in R. Normality was tested in R, using a Shapiro-Wilk's test where the data was small enough, and using Q-Q plots for larger data sets. The Kruskal-Wallis test was used for data not normally to assess if differences between groups were significant. A pairwise Wilcoxon signed-rank test with a Benjamini-Hochburg adjustment

### Supplementary Figures

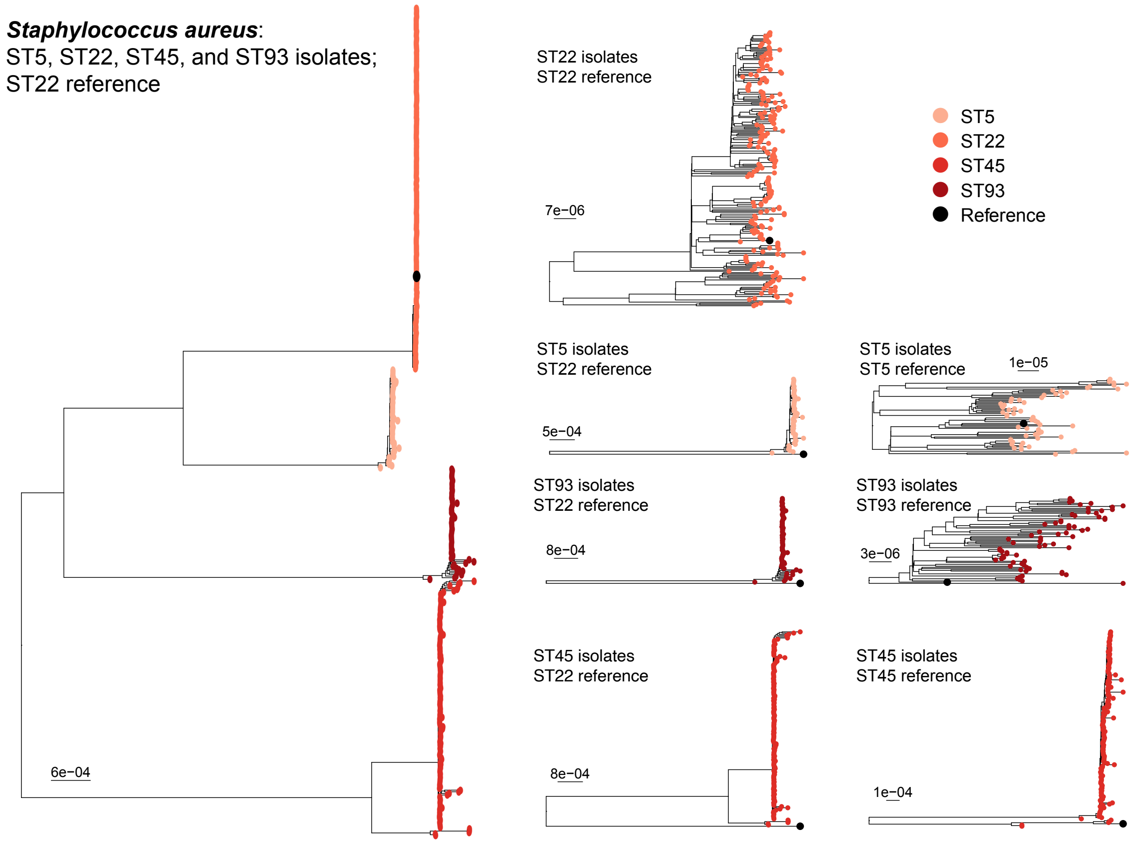


**Supplementary Figure 1. Midpoint-rooted maximum-likelihood phylogenetic trees of *S. aureus* isolates from four sequence types (STs).** Trees include (from left to right): isolates from all four STs and a ST22 reference genome; isolates from within a single ST and a ST22 reference genome, and; isolates from within a single ST and a reference genome of the same ST as the isolates. Tips are coloured by ST.

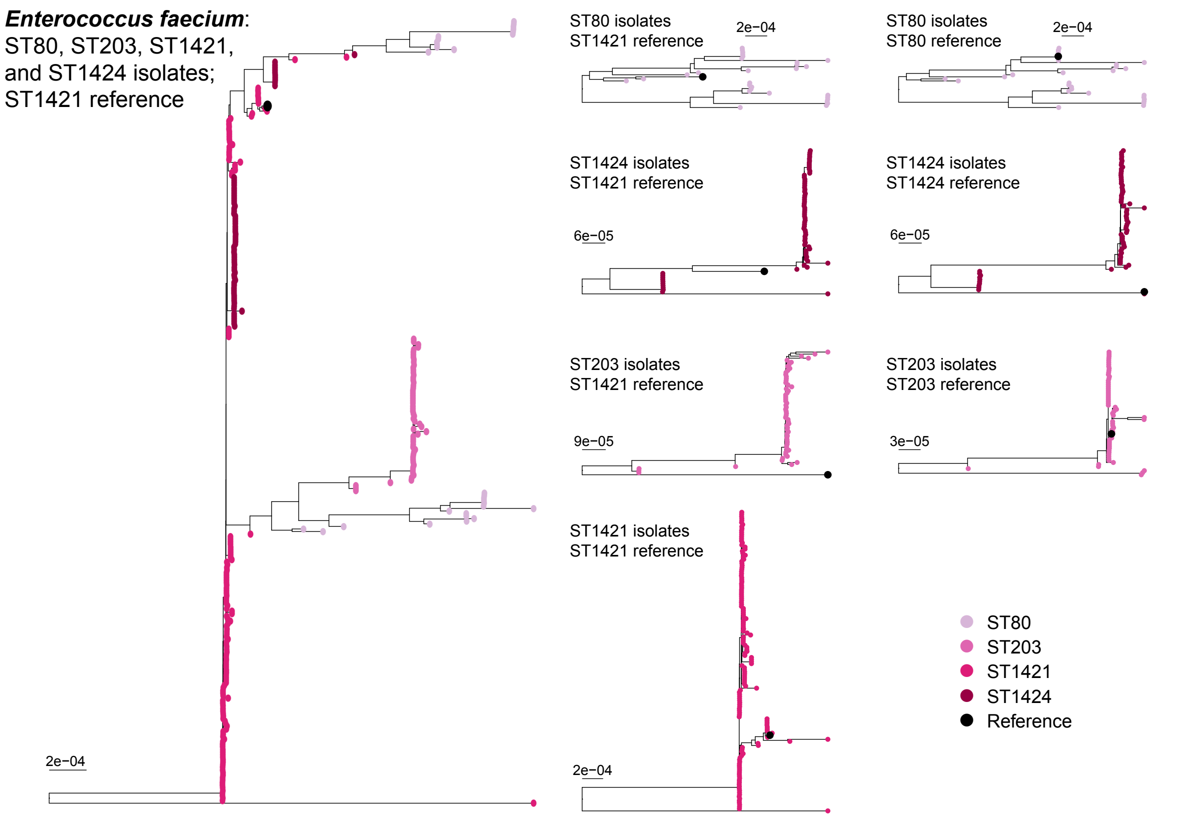


**Supplementary Figure 2. Midpoint-rooted maximum-likelihood phylogenetic trees of *E. faecium* isolates from four sequence types (STs).** Trees include (from left to right): isolates from all four STs and a ST1421 reference genome; isolates from within a single ST and a ST1421 reference genome, and; isolates from within a single ST and a reference genome of the same ST as the isolates. Tips are coloured by ST.

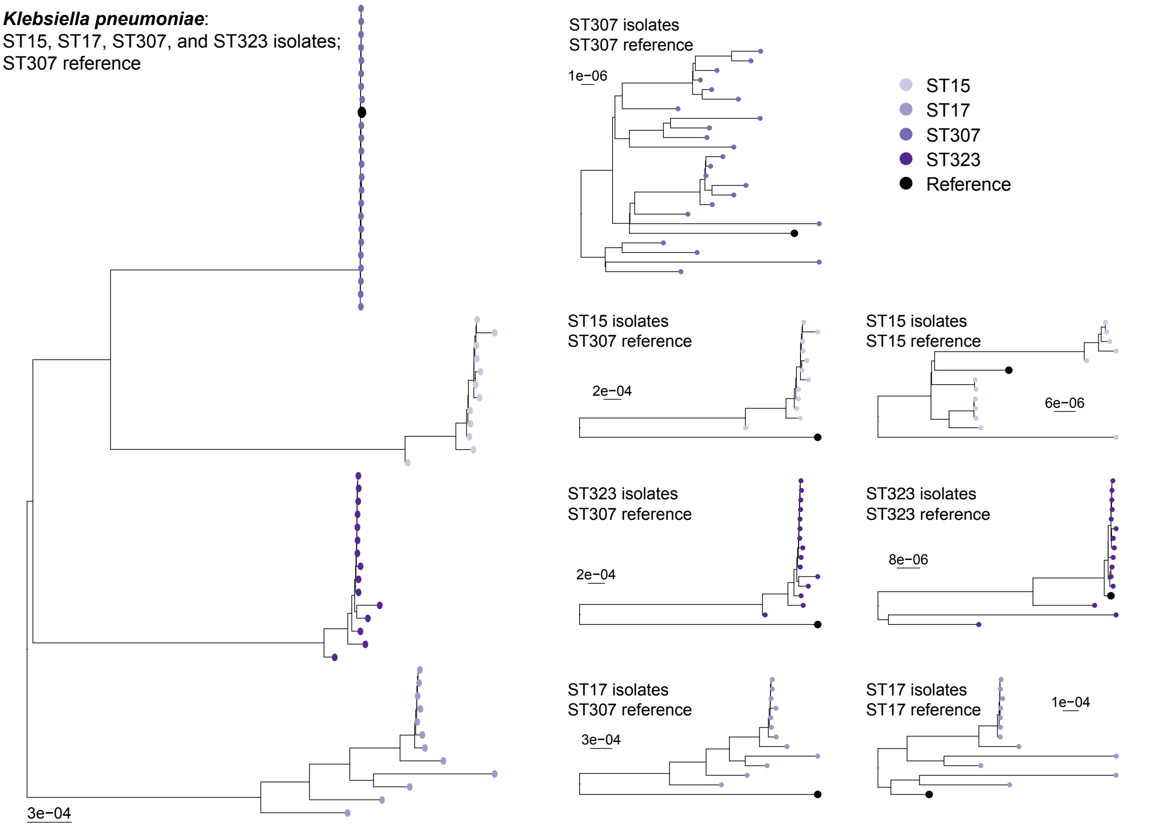


**Supplementary Figure 3. Midpoint-rooted maximum-likelihood phylogenetic trees of *K. pneumoniae* isolates from four sequence types (STs).** Trees include (from left to right): isolates from all four STs and an ST307 reference genome; isolates from within a single ST and an ST307 reference genome, and; isolates from within a single ST and a reference genome of the same ST as the isolates. Tips are coloured by ST.

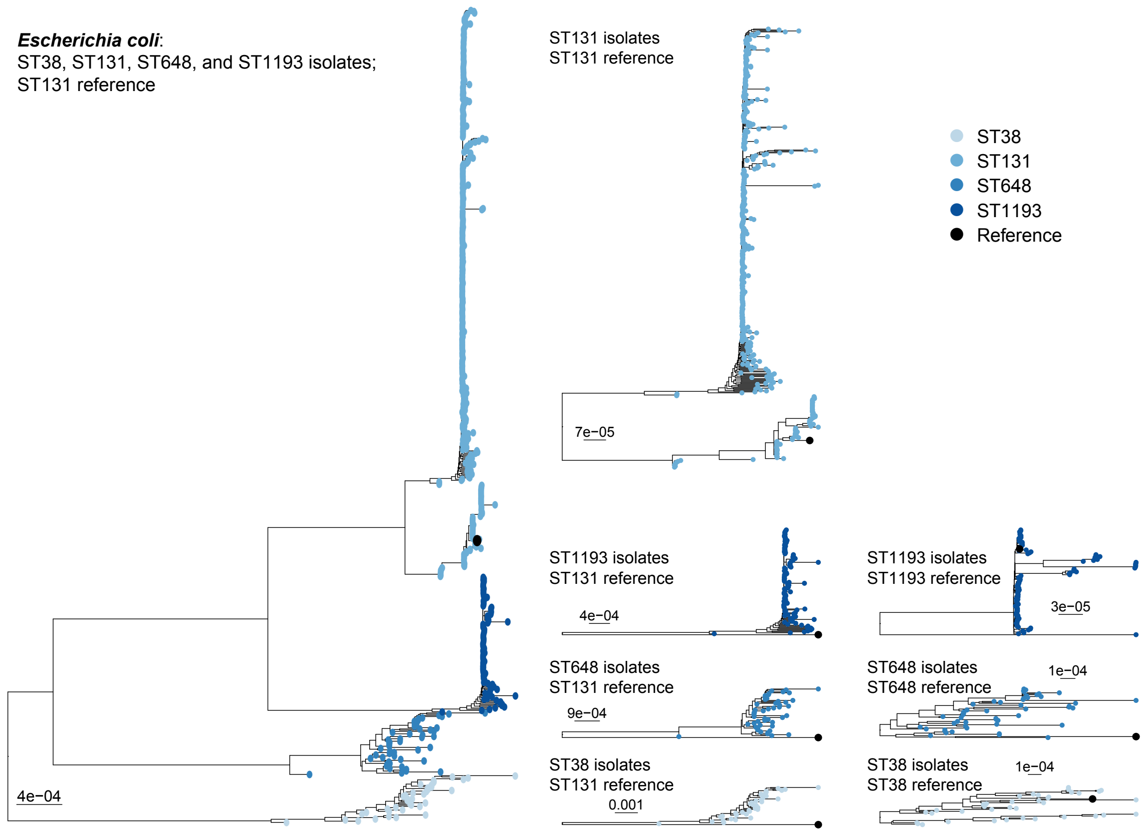


**Supplementary Figure 4. Midpoint-rooted maximum-likelihood phylogenetic trees of *E. coli* isolates from four sequence types (STs).** Trees include (from left to right): isolates from all four STs and an ST38 reference genome; isolates from within a single ST and an ST38 reference genome, and; isolates from within a single ST and a reference genome of the same ST as the isolates. Tips are coloured by ST.

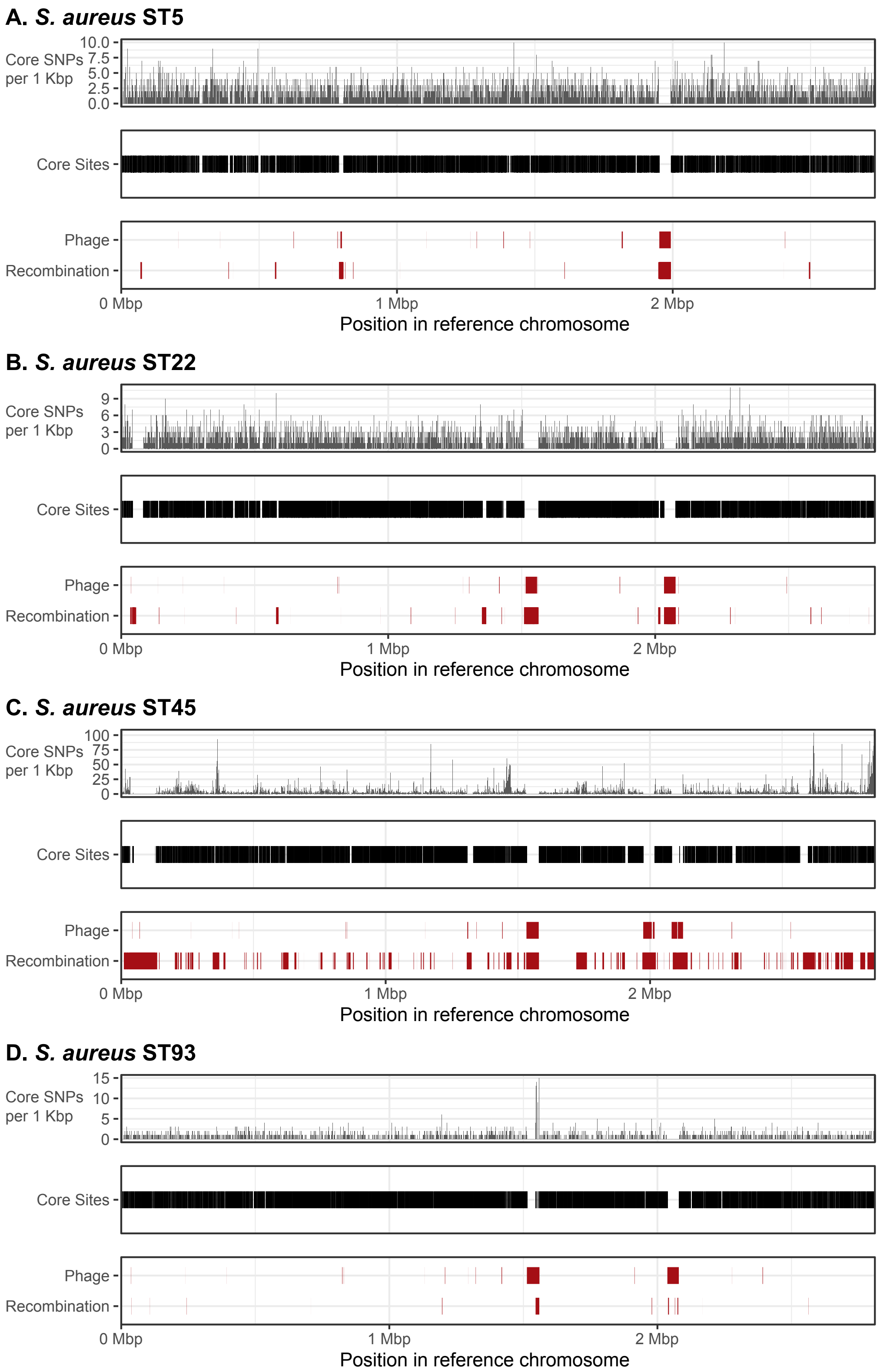


**Supplementary Figure 5. Distribution of core SNPs, core sites, and regions of phage and recombination, and their position in the reference genome for each *S. aureus* sequence type (ST).** Number of core SNPs per 1 Kbp of reference genome length are shown on the y-axis, note the y-axis scale differs depending on ST. Core sites include all sites/base pairs in the reference genome that have an A, T, C, or G nucleotide called in all isolates (i.e. strict core) in the alignment. Phage and recombination regions are shown relative to their position in the reference genome chromosome. The x-axis shows the position and length (maximum x-axis value) of the reference genome chromosome.

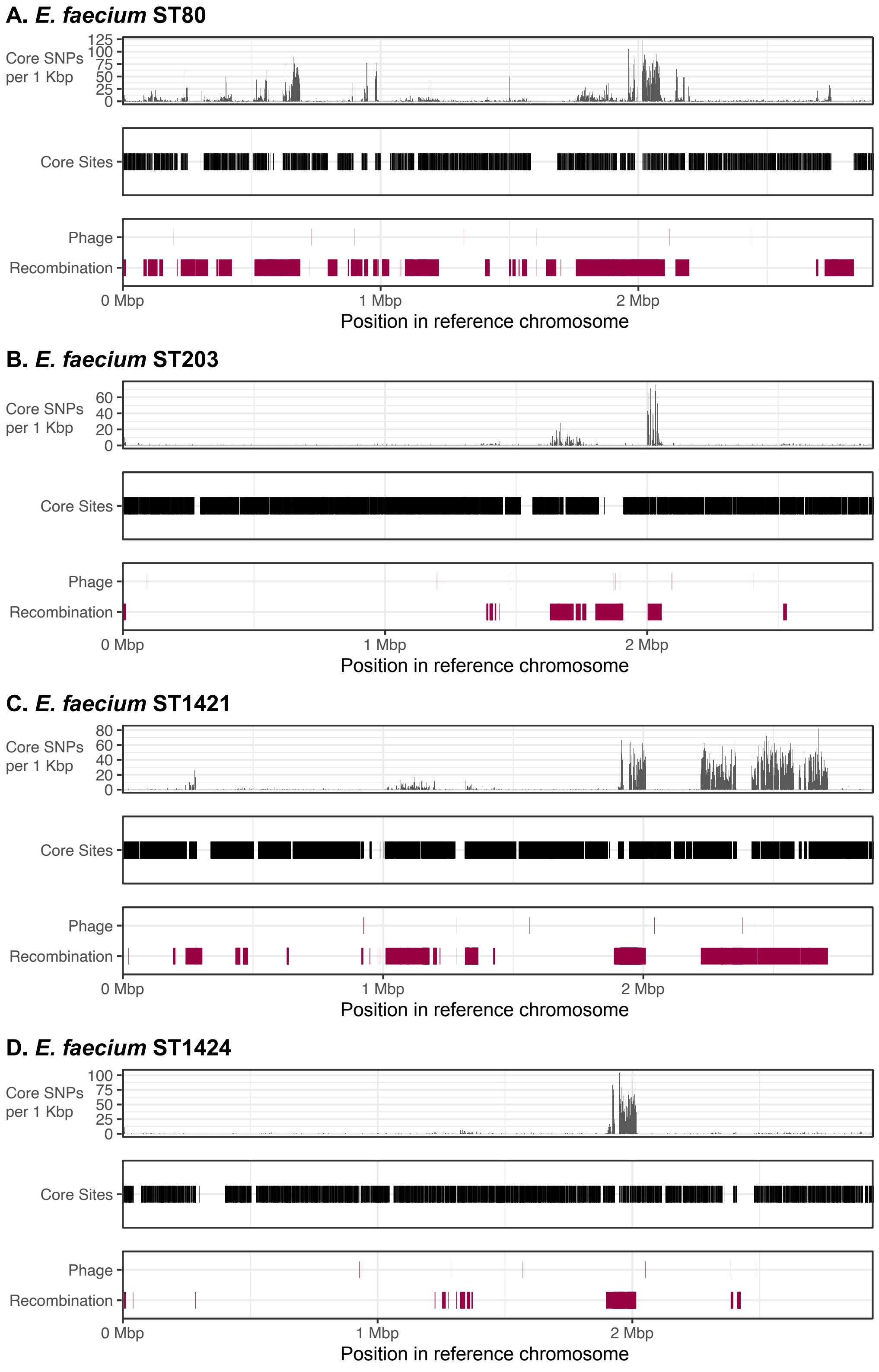


**Supplementary Figure 6. Distribution of core SNPs, core sites, and regions of phage and recombination, and their position in the reference genome for each *E. faecium* sequence type (ST).** Number of core SNPs per 1 Kbp of reference genome length are shown on the y-axis, note the y-axis scale differs depending on ST. Core sites include all sites/base pairs in the reference genome that have an A, T, C, or G nucleotide called in all isolates (i.e. strict core) in the alignment. Phage and recombination regions are shown relative to their position in the reference genome chromosome. The x-axis shows the position and length (maximum x-axis value) of the reference genome chromosome.

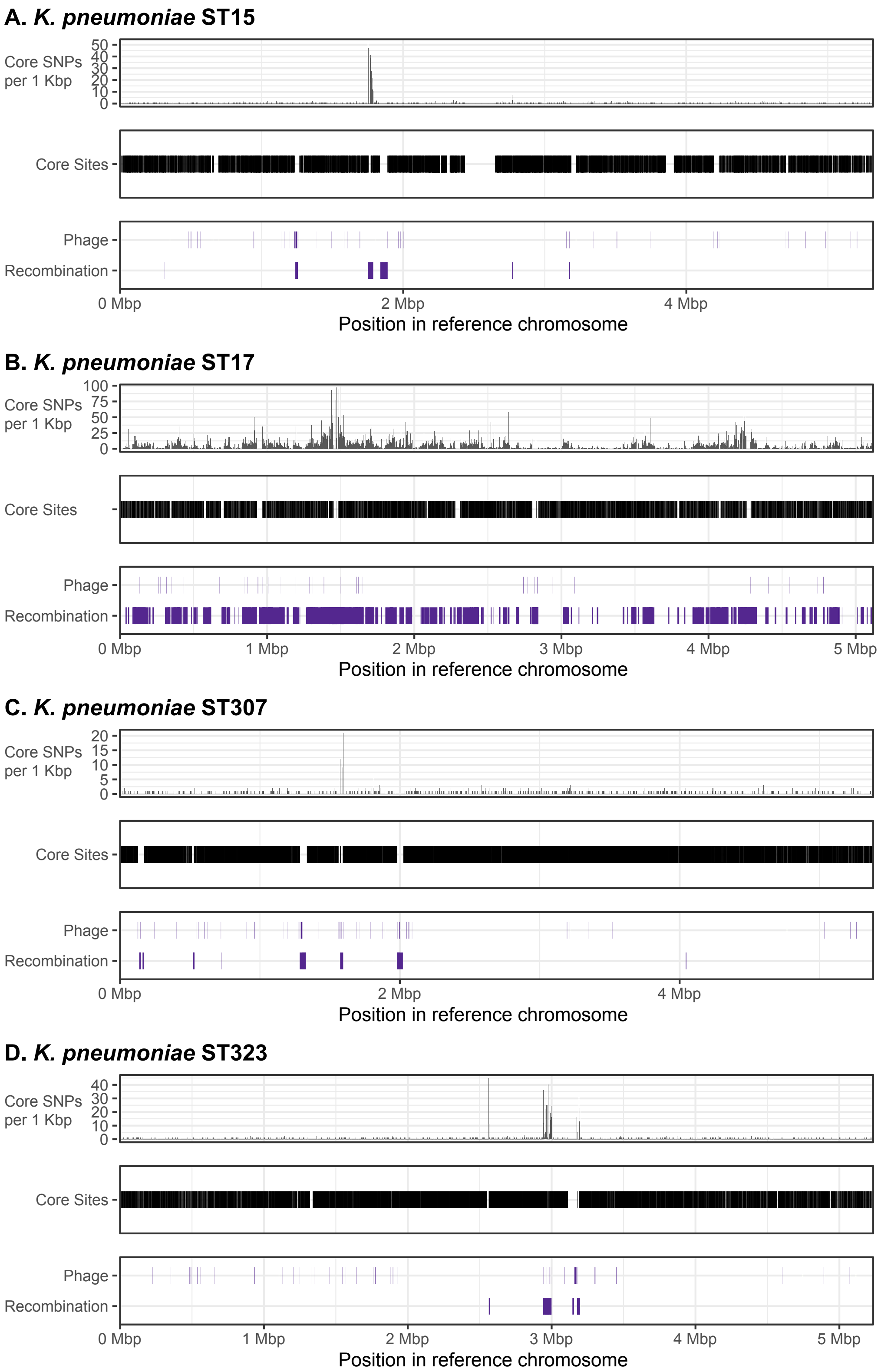


**Supplementary Figure 7. Distribution of core SNPs, core sites, and regions of phage and recombination, and their position in the reference genome for each *K. pneumoniae* sequence type (ST).** Number of core SNPs per 1 Kbp of reference genome length are shown on the y-axis, note the y-axis scale differs depending on ST. Core sites include all sites/base pairs in the reference genome that have an A, T, C, or G nucleotide called in all isolates (i.e. strict core) in the alignment. Phage and recombination regions are shown relative to their position in the reference genome chromosome. The x-axis shows the position and length (maximum x-axis value) of the reference genome chromosome.

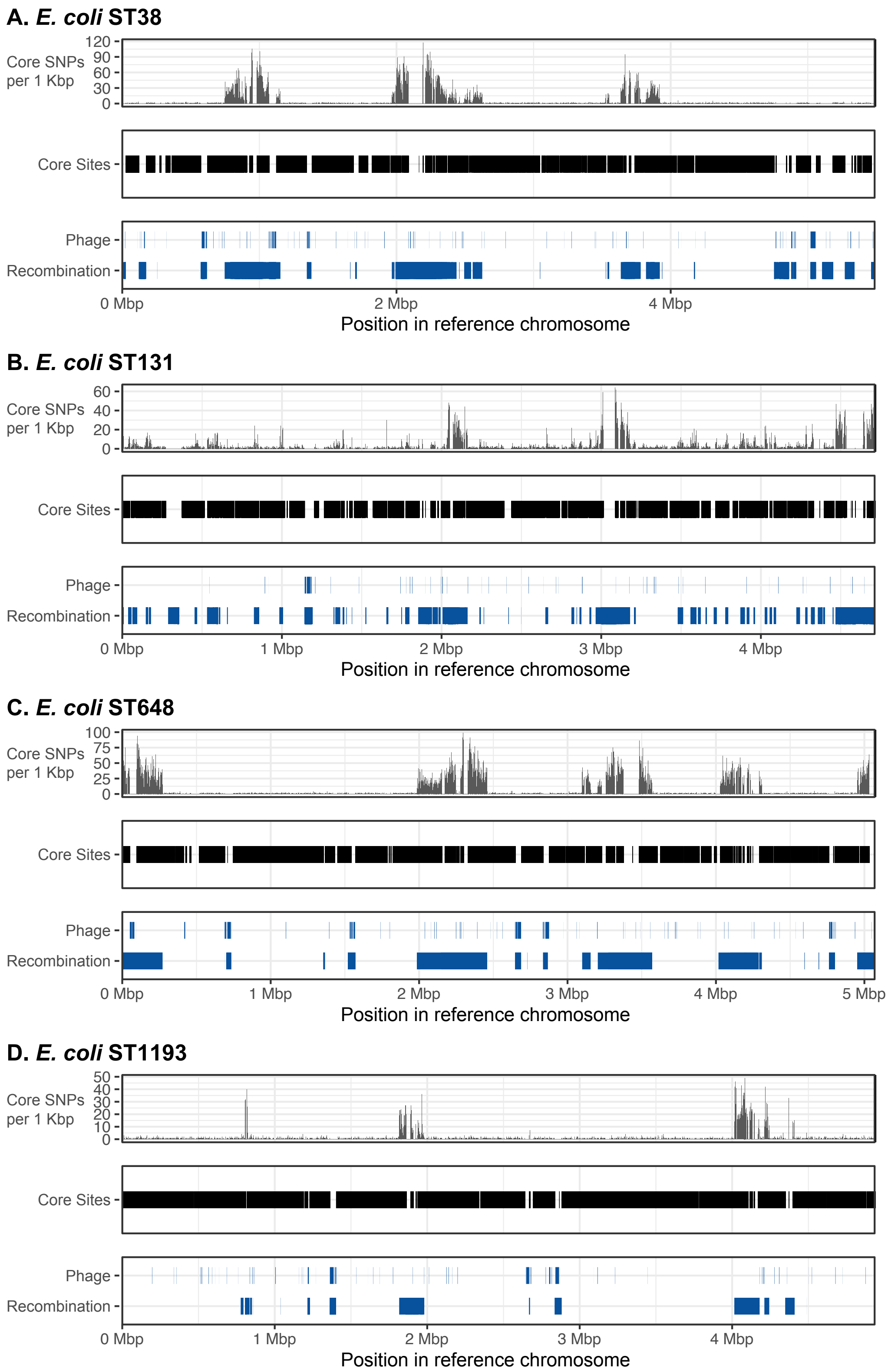


**Supplementary Figure 8. Distribution of core SNPs, core sites, and regions of phage and recombination, and their position in the reference genome for each *E. coli* sequence type (ST).** Number of core SNPs per 1 Kbp of reference genome length are shown on the y-axis, note the y-axis scale differs depending on ST. Core sites include all sites/base pairs in the reference genome that have an A, T, C, or G nucleotide called in all isolates (i.e. strict core) in the alignment. Phage and recombination regions are shown relative to their position in the reference genome chromosome. The x-axis shows the position and length (maximum x-axis value) of the reference genome chromosome.

**Supplementary Table 1. Isolate ID, accession number, species, sequence type (ST), and date of collection (DOC).** [File available on Figshare under DOI: 10.6084/m9.figshare.12992945]

**Supplementary Table 2. Details and results of different analysis levels and reference genome relatedness.** The first column shows the 'Level of Analysis', that is whether this alignment was: 1. Species level (isolates from multiple sequence types (STs) and the chosen reference genome for the species, and per Table 1); 2. Outgroup level (isolates from a single ST but using the same reference genome as the Species level analysis), or; 3. ST level (isolates from a single ST but using a reference genome of the same ST. Information on the species and ST of both the isolates and the reference genome is given in subsequent columns, as well as the total number of isolates of each ST included in the alignment; in Outgroup and ST level analyses this is equivalent to the total number of isolates in the analysis but in the Species level, isolates from all four STs representing the species are included in the alignment. The percentage of the reference genome represented in the core is shown and is calculated from the length of the core alignment (including both variant and invariant sites, given in table) divided by the length of the reference chromosome (also shown). Minimum, maximum and median pairwise SNP distances between i) all isolates of the ST and ii) isolates compared to the reference genome are also provided. The final columns shown the total number of isolate pairs amongst isolates of the same ST, and the number and percentage of isolate pairs falling below the SNP thresholds for putative transmission in each analysis level. [File available on Figshare under DOI: 10.6084/m9.figshare.12992942]

**Supplementary Table 3. Details and results of different prophage and recombination region omission.** The first column shows the 'Level of Analysis', that is whether this was: 1. No masking of any regions; 2. Masking of prophage regions only; 3. Masking of regions of recombination only, or; 4. Masking of both prophage and recombination regions. Information on the species and ST of both the isolates and the reference genome is given in subsequent columns, as well as the total number of isolates of each ST included in the alignment. The percentage of the reference genome represented in the core is shown and is calculated from the length of the core alignment (including both variant and invariant sites, given in table) divided by the length of the reference chromosome (also shown). Minimum, maximum and median pairwise SNP distances between i) all isolates of the ST and ii) isolates compared to the reference genome are also provided. The final columns shown the total number of isolate pairs amongst isolates of the same ST, and the number and percentage of isolate pairs falling below the SNP thresholds for putative transmission in each analysis level. [File available on Figshare under DOI: 10.6084/m9.figshare.12992936"]

**Supplementary Table 4. Details and results of the cumulative approach to isolate inclusion.** Information on the species and ST of both the isolates and the reference genome is given, as well as the total number of isolates of each ST included in the alignment. The maximum date for isolate inclusion for each time period of analysis is shown; all isolates prior to and including this date are included. The percentage of the reference genome represented in the core is shown and is calculated from the length of the core alignment (including both variant and invariant sites, given in table) divided by the length of the reference chromosome (also shown). Minimum, maximum and median pairwise SNP distances between i) all isolates of the ST and ii) isolates compared to the reference genome are also provided. The final columns shown the total number of isolate pairs amongst isolates of the same ST, and the number and percentage of isolate pairs falling below the SNP thresholds for putative transmission in each analysis. [File available on Figshare under DOI: 10.6084/m9.figshare.12992948]

**Supplementary Table 5. Details and results of the sliding window approach to isolate inclusion.** Information on the species and ST of both the isolates and the reference genome is given, as well as the total number of isolates of each ST included in the alignment. The minimum and maximum dates for isolate inclusion for each time window of analysis is shown. The percentage of the reference genome represented in the core is shown and is calculated from the length of the core alignment (including both variant and invariant sites, given in table) divided by the length of the reference chromosome (also shown). Minimum, maximum and median pairwise SNP distances between i) all isolates of the ST and ii) isolates compared to the reference genome are also provided. The final columns shown the total number of isolate pairs amongst isolates of the same ST, and the number and percentage of isolate pairs falling below the SNP thresholds for putative transmission in each analysis. [File available on Figshare under DOI: 10.6084/m9.figshare.12992951]
